# Redundant γc cytokines license IL-1-driven neutrophil inflammation through MEK/ERK convergence

**DOI:** 10.64898/2026.05.28.728506

**Authors:** Kristian Lorenzo, Lauren Arayan, Timothy M. Stearns, Lisa M. Burzenski, Jing Wen, Leonard D. Shultz, Vishnu Hosur

## Abstract

Interleukin-1 (IL-1) is a central driver of autoinflammatory disease, yet IL-1 blockade often provides incomplete benefit in complex, neutrophil-driven conditions. Here we identify a licensing circuit in which common γ-chain (γc) cytokines provide a redundant signal required for maximal IL-1–driven neutrophil inflammation. IL-1 and γc cytokines synergize to drive inflammatory cytokine production exceeding either stimulus alone, and these signals engage the MEK/ERK pathway, an effect substantially suppressed by pharmacological MEK inhibition. We validated this circuit in vivo in a mouse model of IL-1α–driven neutrophil-dominant autoinflammation. Ablation of the shared γc receptor markedly prolonged survival and attenuated pathology, whereas deletion of individual γc cytokine pathways had no major effect—demonstrating in vivo necessity and functional redundancy. Analysis of public phospho-proteomic and transcriptomic datasets confirms MEK/ERK as a conserved neutrophil response to diverse inflammatory stimuli and coordinated IL-1, γc, and MEK/ERK activation in neutrophils from patients with systemic juvenile idiopathic arthritis (sJIA) and in lesional skin from hidradenitis suppurativa. Together, these findings define a signaling architecture in which redundant γc inputs enhance MEK/ERK-dependent inflammatory output, identify the γc receptor as an in vivo disease-modifying node, and position MEK/ERK as a mechanistically grounded therapeutic target.

**eTOC Summary:** Lorenzo et al. show that common γ-chain (γc) cytokines provide redundant licensing signals that amplify IL-1–driven neutrophil inflammation through MEK/ERK convergence. Blocking any single γc cytokine fails to suppress disease, but ablating the shared γc receptor or inhibiting MEK/ERK markedly attenuates pathology, identifying these nodes as therapeutic targets in autoinflammatory disease.

## INTRODUCTION

Interleukin-1 (IL-1) is among the most potent orchestrators of innate immunity, yet its therapeutic blockade produces paradoxically inconsistent results across inflammatory diseases. In monogenic syndromes such as cryopyrin-associated periodic syndromes (CAPS), IL-1 inhibition is transformative (*1, 2*). Yet in neutrophil-dominant conditions—including systemic juvenile idiopathic arthritis (sJIA) and adult-onset Still’s disease (AOSD)—the same strategy yields only partial or inconsistent benefit (*2, 3*). This clinical divergence raises a fundamental question of inflammatory gain control: what determines whether IL-1 signaling produces limited, self-resolving activation versus destructive, high-output inflammation? We propose that the answer lies not within the IL-1 pathway itself, but in a parallel licensing mechanism that sets the inflammatory gain of the IL-1 response.

A compelling candidate for this licensing function resides in the common γ-chain (γc; IL2RG) cytokine family4, which includes IL-2, IL-4, IL-7, IL-9, IL-15, and IL-21. Although canonically studied for their roles in lymphocyte development and adaptive immunity (*4*), γc cytokine receptors are expressed on innate immune cells, including neutrophils and macrophages(*5, 6*). Signaling through these receptors in myeloid cells is actively constrained by intracellular negative regulators, most critically the protein tyrosine phosphatase SHP-1 (encoded by *Ptpn6*). Mechanistically, SHP-1 enforces a quantitative ceiling on cytokine receptor output: its dephosphorylation of JAK-associated receptor complexes sets the gain of downstream JAK–STAT and MAPK signaling, and genetic or pharmacological loss of SHP-1 function is sufficient to breach the threshold for pathological neutrophil activation and drive systemic autoinflammation in mice and humans (*5, 7, 8*).

Direct evidence that γc signaling is engaged in human neutrophil-driven disease comes from re-analysis of two anatomically and etiologically distinct public transcriptomic datasets. In peripheral blood neutrophils from systemic juvenile idiopathic arthritis (sJIA) patients in GSE103170(*9*), and in lesional skin from patients with hidradenitis suppurativa (HS) in GSE148027(*10*), the IL-1, γc, and MEK/ERK signaling axes are coordinately and significantly upregulated relative to healthy controls. Downstream transcriptional targets of IL-1/NFκB and γc/JAK-STAT signaling are concurrently enriched across these two re-analyzed datasets, consistent with cytokine-driven pathway activation(*3, 11-13*). Transcriptional changes in core MEK/ERK pathway component genes are, by contrast, modest—consistent with ERK activity being regulated predominantly at the post-translational level rather than through proportional ligand-driven gene induction. This pattern of pathway activation without commensurate transcriptional upregulation of its components is characteristic of paracrine cytokine signaling and positions γc input as a modulator of IL-1 responsiveness, rather than an independent inflammatory driver.

These observations raise the question of how this paracrine input is transduced into amplified inflammatory output. We hypothesize that γc cytokines provide interchangeable licensing signals that synergize with IL-1 to amplify neutrophil inflammatory output. Because individual γc family members signal through the shared γc receptor subunit and overlapping downstream JAK/STAT machinery(*6*), loss of any single ligand would be predicted to leave the circuit largely intact. This logic helps explain why inhibition of a single γc cytokine—or of IL-1 alone—may fail to fully suppress inflammation when parallel γc input persists. At the same time, the amplified response should depend on downstream signaling that can be engaged by both inputs. Based on the transcriptional and pharmacological evidence presented here, MEK/ERK emerges as the strongest candidate for that shared pathway.

To dissect this circuit, we combine *in vitro* cell biology, unbiased transcriptomics, and targeted pharmacological perturbation in primary mouse and human neutrophils with a powerful *in vivo* genetic system. We exploit a spontaneous loss-of-function mutation in *Ptpn6*—designated *me4J*—that causes lethal, neutrophil-dominant autoinflammation driven by IL-1α signaling in NOD *scid* mice(*8, 14*). This model, in which SHP-1 deficiency creates a sensitized signaling context, enabled us to test the *in vivo* necessity and sufficiency of individual components of the licensing circuit through systematic genetic crosses.

We show that γc cytokines and IL-1α synergize to drive inflammatory cytokine production, and that this synergy engages MEK/ERK signaling. *In vivo*, while individual γc cytokines are fully dispensable—confirming functional redundancy—the shared γc receptor chain is required for full neutrophil-driven inflammatory disease, while downstream MEK/ERK signaling emerges as the likely common effector pathway supported by *in vitro* pharmacology and signaling analysis. Together, these findings define a signaling architecture that governs IL-1–driven neutrophil inflammation and provide a principled explanation for why single-pathway blockade so frequently fails in autoinflammatory disease.

## RESULTS

### A coordinated IL-1/γc/MEK-ERK signaling axis is a conserved feature of human autoinflammatory disease

To assess the clinical relevance of the IL-1, common γ-chain (γc), and MEK/ERK signaling axes, we analyzed publicly available transcriptomic data from two distinct neutrophil-driven autoinflammatory diseases. In peripheral blood neutrophils from patients with sJIA in GSE103170(*9*), gene sets corresponding to IL-1, γc, and MEK/ERK pathway components were significantly and coordinately upregulated relative to healthy controls (Figure 1A), as corroborated by pathway module scores and Gene Set Enrichment Analysis (GSEA) (Figures 1B and 1C). Downstream transcriptional targets of IL-1/NF-κB and γc/JAK-STAT signaling, together with convergent MEK/ERK outputs, were likewise significantly enriched in disease (Figures 1D–1F). Notably, transcriptional changes in core MEK/ERK pathway component genes were modest in Figure 1A, consistent with ERK activity being regulated predominantly at the post-translational level. We confirmed and extended these findings in a second, anatomically distinct cohort: bulk skin transcriptomes from patients with hidradenitis suppurativa (HS) in GSE148027(*10*) revealed a similarly coordinated pattern of IL-1/γc/MEK-ERK pathway activation across lesional versus non-lesional comparisons (Figures 1A–1F). Together, these data establish that co-activation of the IL-1, γc, and MEK/ERK axes is a conserved transcriptional feature of human autoinflammatory disease (see Supplemental Table 1 for full gene lists).

**Fig. 1.**
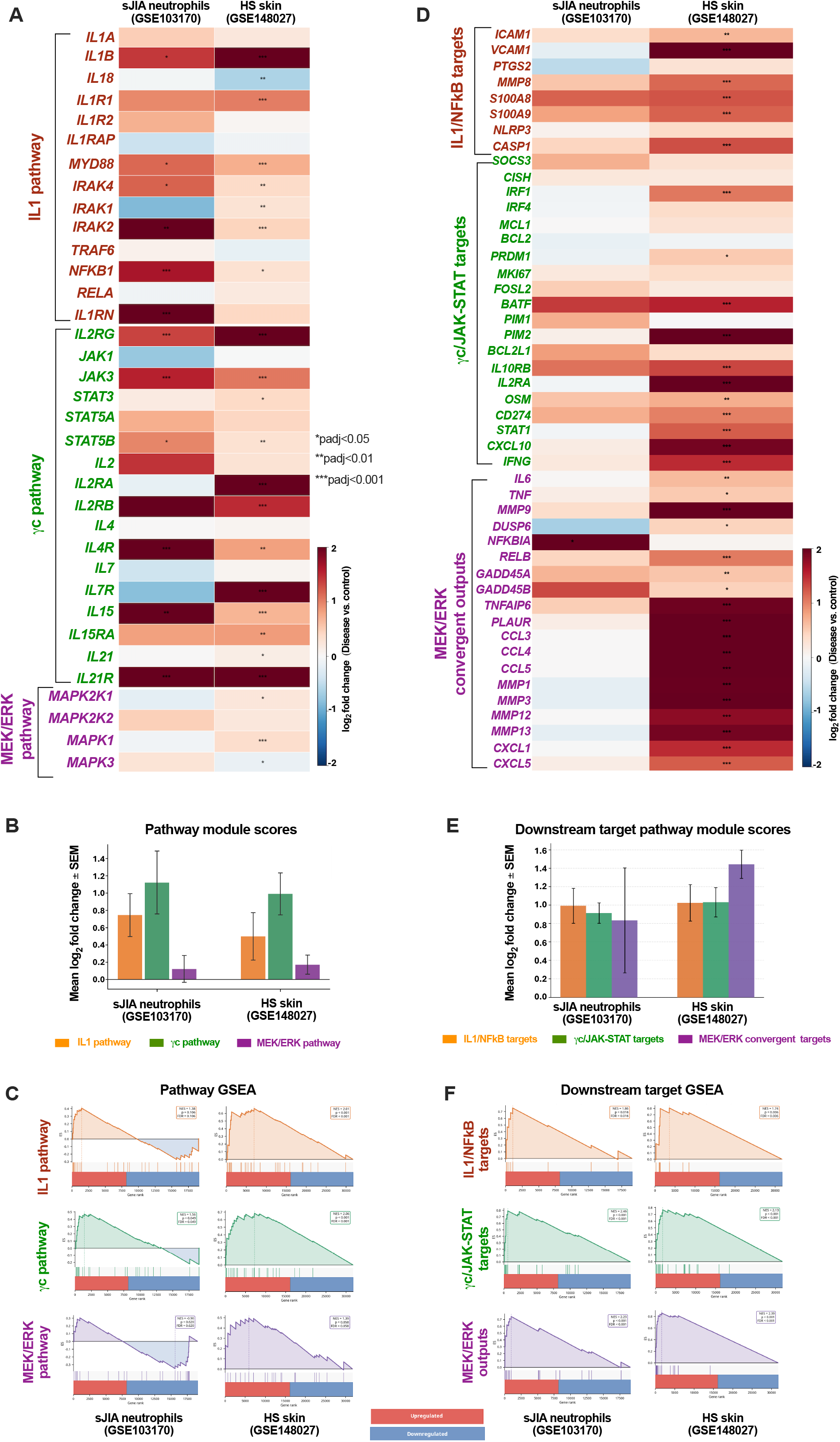
Coordinated upregulation of IL-1, γc, and MEK/ERK signaling pathways in human autoinflammatory disease. To determine the clinical relevance of the IL-1, common γ-chain (γc), and MEK/ERK signaling axes, we analyzed publicly available transcriptomic data from two distinct neutrophil-driven autoinflammatory diseases: systemic juvenile idiopathic arthritis (sJIA; peripheral blood neutrophils; GSE103170; n=3/group) and hidradenitis suppurativa (HS; lesional skin biopsies; GSE148027; n=18 lesional, n=15 non-lesional/healthy). (A and D) Heatmaps showing the log_2_ fold change (Disease vs. Control) of curated gene sets for upstream pathway components (A) or downstream transcriptional targets (D). Gene names are color-coded by pathway module: red, IL-1 pathway; green, γc pathway; purple, MEK/ERK pathway. Asterisks denote Benjamini-Hochberg-adjusted p values (*p < 0.05, **p < 0.01, ***p < 0.001). (B and E) Pathway module scores for upstream pathways (B) or downstream target gene sets (E), calculated as the mean log_2_ fold change of significantly differentially expressed genes within each gene set. Bars represent mean ± SEM. (C and F) Gene Set Enrichment Analysis (GSEA) of upstream pathways (C) or downstream target gene sets (F), performed on gene lists ranked by t-statistic from differential expression analysis. NES, normalized enrichment score; FDR, false discovery rate.

### Redundant γc cytokines license IL-1-driven neutrophil inflammation and converge on MEK/ERK signaling

Co-stimulation of NOD *scid* +/+ bone marrow–derived neutrophils with IL-1α or IL-1β and the γc cytokine IL-15 produced synergistic amplification of IL-6 and TNFα secretion, with a pronounced leftward shift in the IL-1 dose–response curve indicating that γc co-stimulation dramatically lowers the threshold for IL-1– driven cytokine production (**Figures 2A–2D**). IL-2 co-stimulation produced equivalent amplification of IL-1– driven IL-6 secretion (**Figures 2E and 2F**), establishing that this licensing effect is not restricted to a single γc family member. IL-4 and IL-7 each recapitulated the synergistic amplification observed with IL-2 and IL-15 (**Figures 2G and 2H**), whereas IL-9 and IL-21 failed to enhance IL-1α–induced IL-6 production (**Figures 2I and 2J**; p = 0.06 and p = 0.7, respectively), demonstrating that licensing capacity is a shared but not universal property of the γc cytokine family.

**Fig. 2.**
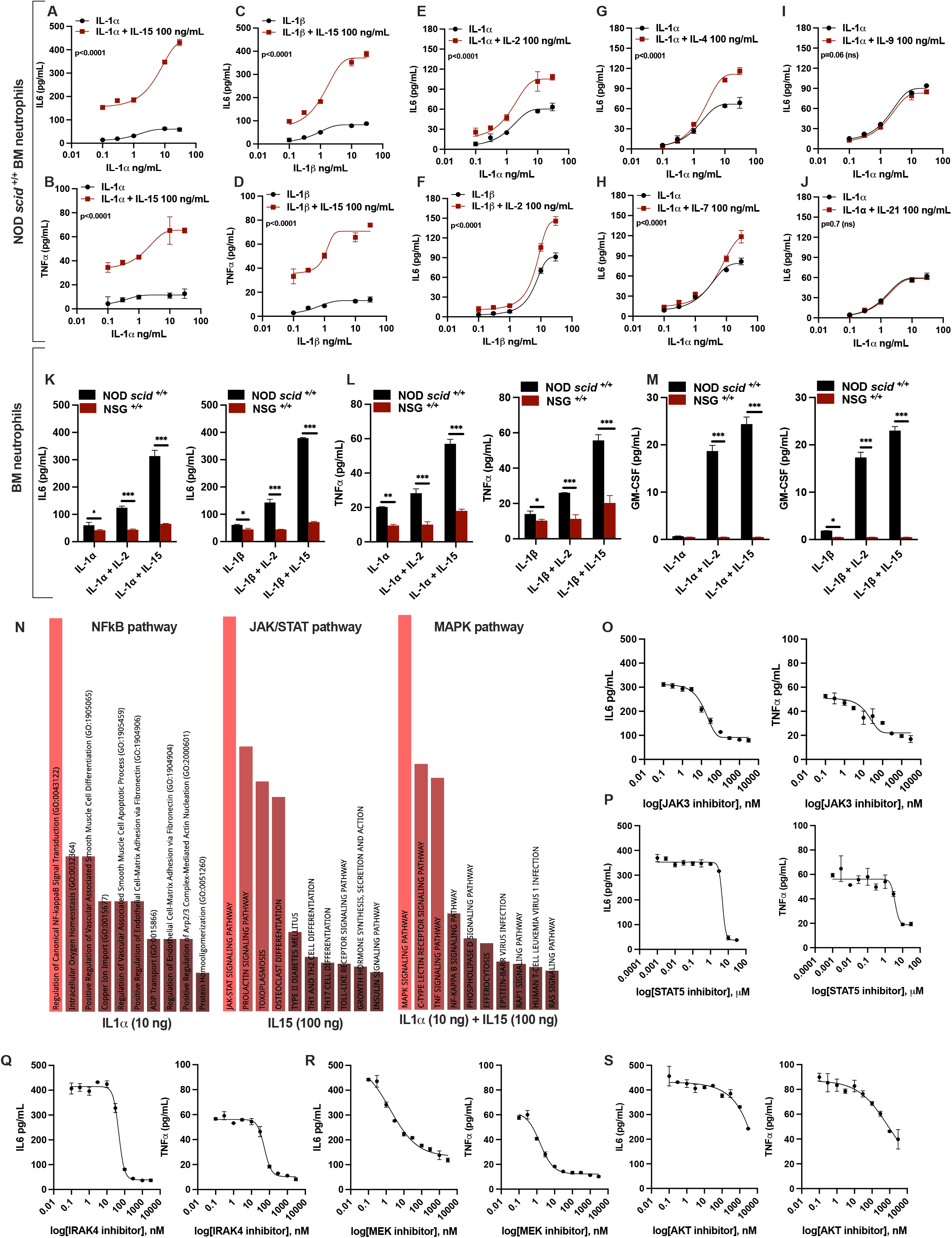
Common γ-chain cytokines license IL-1-driven inflammatory cytokine production in neutrophils and activate MEK/ERK signaling. (A–J) Dose-response curves for IL-6 secretion (A–C, E–J) or TNFα secretion (B, D) by NOD scid +/+ bone marrow neutrophils stimulated with increasing concentrations of IL-1α (A, B, E, G–J) or IL-1β (C, D, F) alone (black circles) or in combination with a fixed concentration (100 ng/mL) of the indicated γc cytokine (red squares): IL-15 (A–D), IL-2 (E, F), IL-4 (G), IL-7 (H), IL-9 (I), or IL-21 (J). Data are mean ± SEM; n=3 biological samples, each measured in duplicate by ELISA. Statistical significance determined by two-way ANOVA with multiple comparisons. (K–M) IL-6 (K), TNFα (L), and GM-CSF (M) secretion by BM neutrophils from NOD scid +/+ (black) or NSG +/+ (red) mice stimulated with IL-1α (10 ng/mL) or IL-1β (10 ng/mL) alone or in combination with IL-2 (100 ng/mL) or IL-15 (100 ng/mL) for 18 h. Data are mean ± SEM; n=3 biological samples per genotype. *p < 0.05, **p < 0.01, ***p < 0.001 by one-way ANOVA with Tukey’s multiple comparisons test. (N) Gene Ontology (GO) biological process enrichment analysis of RNA-sequencing data from NOD scid +/+ BM neutrophils stimulated for 3 h with IL-1α (10 ng/mL), IL-15 (100 ng/mL), or IL-1α + IL-15 in combination. Top enriched GO terms are ranked by −log10 p value. (O–S) Dose-dependent inhibition of IL-6 (left) and TNFα (right) secretion by NOD scid +/+ BM neutrophils co-stimulated with IL-1α (30 ng/mL) + IL-15 (100 ng/mL) in the presence of increasing concentrations of the JAK3 inhibitor ritlecitinib (O), the STAT5 inhibitor AC-4-130 (P), the IRAK4 inhibitor zimlovisertib (Q), the MEK inhibitor trametinib (R), or the AKT inhibitor capivasertib (AZD5363) (S). Data are mean ± SEM; n=3 biological samples, each measured in duplicate by ELISA.

The functional redundancy among IL-2, IL-4, IL-7, and IL-15 implicated the shared γc receptor chain (IL2RG) as the critical upstream node. Synergistic amplification of IL-6, TNFα, and GM-CSF secretion induced by IL-2 and IL-15 co-stimulation was abrogated in γc-deficient NSG +/+ bone marrow neutrophils (**Figures 2K– 2M**), establishing that γc receptor signaling is required for the licensing effect.

To define the transcriptional architecture integrating IL-1 and γc inputs, RNA sequencing was performed on neutrophils stimulated with IL-1α, IL-15, or both in combination. IL-1α alone induced canonical NFκB target genes; IL-15 alone preferentially activated JAK–STAT targets. Co-stimulation produced a transcriptional program exceeding the sum of individual stimuli, with Gene Ontology enrichment analysis of synergistically induced genes revealing selective enrichment of MAPK and MEK/ERK signaling pathways (**Figure 2N**). Pathway-specific gene expression patterns for NFκB, JAK/STAT, and MAPK target genes across all three stimulation conditions are detailed in Supplemental Figure 1.

Pharmacological dissection identified MEK/ERK as the dominant downstream pathway under these conditions. MEK inhibition with trametinib produced substantial suppression of both IL-6 and TNFα across the full dose range (Figure 2R). The IRAK4 inhibitor zimlovisertib likewise fully suppressed cytokine output (Figure 2Q), confirming obligate IL-1 receptor engagement. JAK3 and STAT5 inhibition produced dose-dependent but incomplete suppression (Figures 2O and 2P), consistent with γc–JAK/STAT signaling amplifying rather than independently driving inflammatory output. The AKT inhibitor capivasertib (AZD5363) produced only a partial reduction in IL-6 and TNFα production, and only at supra-pharmacological concentrations (>1 μM) that likely reflect off-target effects rather than on-target AKT inhibition (Figure 2S), suggesting that AKT is not a primary mediator of the IL-1+IL-15 licensing circuit under these conditions.

### IL-1 and γc signaling engage MEK/ERK in mouse and human neutrophils

To directly test whether ERK functions as a biochemical convergence point, pERK kinetics were measured in mouse neutrophils by phospho-flow cytometry. Although IL-1α or IL-15 alone induced modest ERK phosphorylation, co-stimulation produced rapid, synergistic amplification of pERK evident in the frequency of responding cells across three independent dose conditions (see also Supplemental Figure 2; **Figures 3A–F**), as well as in per-cell signal intensity at the highest dose (**Figure 3G**). This synergistic pERK response was abrogated by the MEK inhibitor trametinib (**Figure 3H**), confirming that the convergent signal is MEK-dependent.

**Fig. 3.**
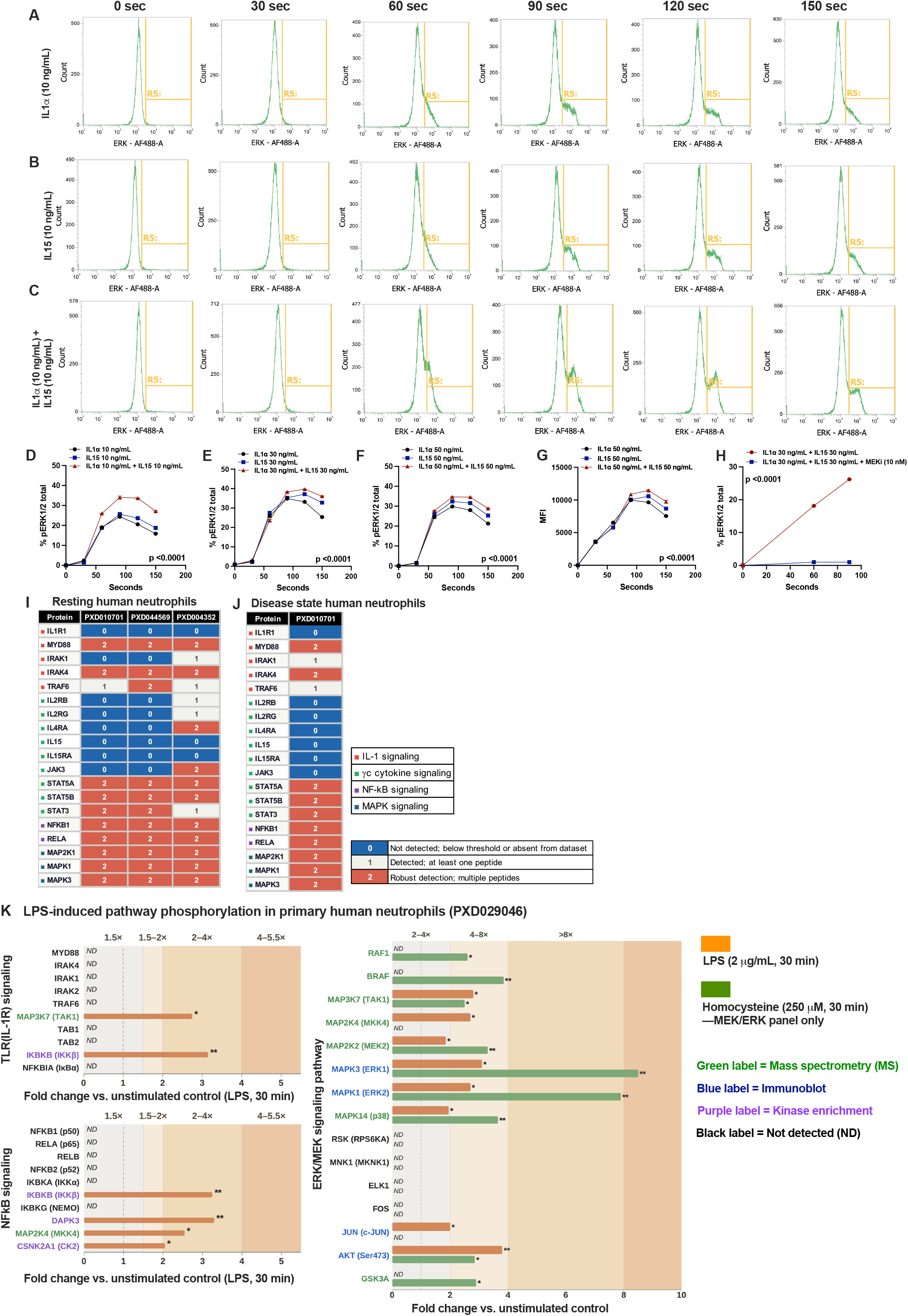
MEK/ERK activation is a conserved feature of inflammatory signaling in mouse and human neutrophils. To test whether this pathway behavior is conserved in human cells, we first used a mouse model to define the kinetic and synergistic signaling dynamics (A–H), then confirmed the presence and activity of the core pathway components in primary human neutrophils through a cross-study analysis of public proteomic and phospho-proteomic data (I–K). **(A–H) Synergistic ERK phosphorylation in mouse neutrophils: (A–C)** Representative intracellular flow cytometry histograms showing phospho-ERK1/2 (pERK) in NOD *scid* +/+ bone marrow neutrophils stimulated with IL-1α (10 ng/mL; A), IL-15 (10 ng/mL; B), or both (C) over 150 seconds. The R5 gate (yellow) demarcates the pERK-positive population. **(D–E)** Quantification of the percentage of pERK1/2-positive cells over time for the 10 ng/mL (D) and 30 ng/mL (E) stimulation conditions. Co-stimulation (IL-1α + IL-15) produced a synergistic and sustained pERK response that was significantly greater than either stimulus alone in the proportion of responding cells. **(F–G)** Quantification of the percentage of pERK1/2-positive cells (F) and pERK median fluorescence intensity (MFI; G) at 50 ng/mL, demonstrating that synergistic ERK amplification is dose-independent and increases both the proportion of responding cells and per-cell signal intensity. **(H)** Pre-treatment with the MEK inhibitor trametinib (10 nM) completely abrogated the synergistic pERK response, confirming that signal convergence on ERK is entirely MEK-dependent. Data in (D–H) are mean ± SEM; *n*=3 biological replicates. \*\*\**p* < 0.001 by two-way ANOVA. **(I–K) Proteomic and phospho-proteomic evidence in human neutrophils: (I–J)** Detection of core IL-1, γc, and MEK/ERK pathway proteins across independent public human neutrophil proteomics datasets from resting healthy donor neutrophils (I; PXD010701, PXD044569, PXD004352) and disease-state neutrophils (J; PXD010701). Each cell reflects detection confidence per dataset, scored as follows: 0, not detected or below threshold; 1, detected with at least one peptide; 2, robust detection with multiple peptides. **(K)** Stimulus-induced phosphorylation changes for pathway components compiled from public phospho-proteomic datasets (PXD029046) and PhosphoSitePlus annotations. Bar length represents the magnitude of phosphorylation change relative to unstimulated controls; black bars indicate dephosphorylation. LPS, a TLR4 agonist sharing the IL-1R signaling axis, induces comparatively modest ERK phosphorylation relative to homocysteine, consistent with the requirement for co-stimulatory input to achieve full MEK/ERK convergence observed in mouse neutrophils.

To establish whether this signaling behavior is conserved in human neutrophils, core components of the IL-1, γc, and MEK/ERK pathways were detected across independent public proteomic datasets from both healthy donors and patients with inflammatory disease (**Figures 3I and 3J**). Phospho-proteomic analysis of primary human neutrophils(*15*) further demonstrated that LPS—a TLR4 agonist that shares the MyD88/IRAK/TRAF6 signaling axis with IL-1R—induced only modest ERK phosphorylation, whereas the thrombosis-associated metabolic signal homocysteine induced substantially stronger ERK activation (**Figure 3K**). This divergence in ERK output mirrors the weak pERK response to IL-1α alone observed in mouse neutrophils and supports the idea that TLR4/IL-1R-type stimulation is an intrinsically modest driver of ERK activation in human neutrophils unless additional input is present. Together, these proteomic and phospho-proteomic data establish that the core IL-1, γc, and MEK/ERK pathway components are expressed and activatable in human neutrophils, supporting conservation of this signaling architecture across species (**Figures 3I–K**).

### A spontaneous loss-of-function mutation in *Ptpn6* causes lethal neutrophil-dominant autoinflammation in NOD *scid* mice

To investigate whether dysregulated cytokine receptor signaling in neutrophils is sufficient to drive autoinflammation *in vivo*, we sought a genetically tractable model of myeloid-intrinsic signaling hyperactivation. SHP-1 (encoded by *Ptpn6*) acts as a critical phosphatase brake on cytokine receptor signaling in myeloid cells, and its loss de-represses these pathways, providing a sensitized signaling context in which inflammatory circuits become pathologically amplified. During routine colony maintenance, we identified a spontaneous recessive mutation in our NOD *scid* colony—designated motheaten-4J (*me4J*)—that produced a severe systemic inflammatory phenotype (**Figures 4A and 4B**). Homozygous NOD *scid*-*me*^*4J/*^*me*^*4J*^ mice displayed profound growth retardation, severe ulcerative skin lesions, reduced body weight, and elevated footpad inflammation scores compared with heterozygous +/*me*^*4J*^ littermates (**Figures 4C, 4D**). Survival was markedly curtailed in both sexes, with no significant difference in disease course between males and females (**Figure 4E**; p = ns).

**Fig. 4.**
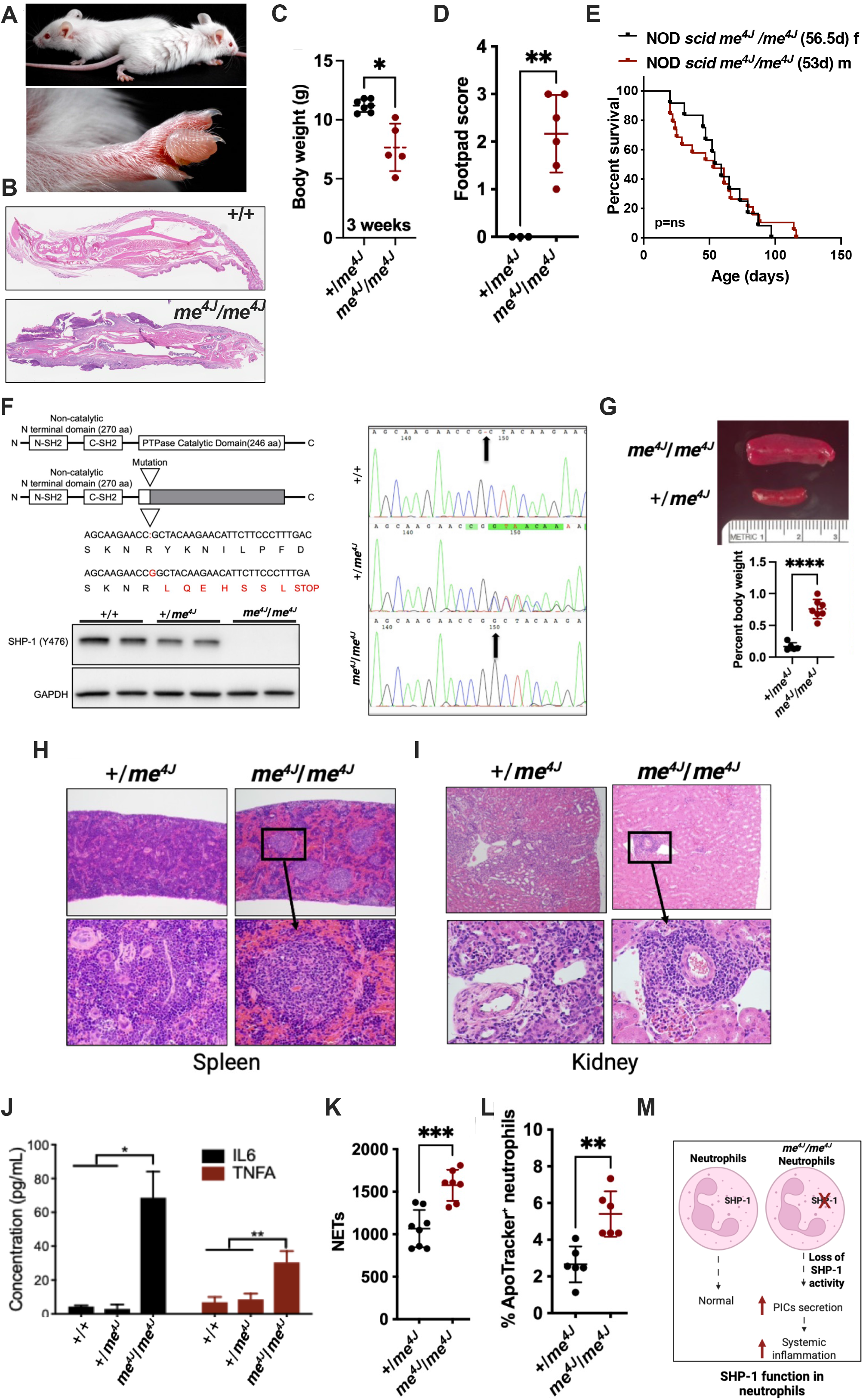
A spontaneous loss-of-function mutation in *Ptpn6* (SHP-1) in the NOD *scid* background causes severe neutrophil-driven autoinflammatory disease. **(A)** Gross phenotype of NOD *scid*-*me*^*4J*^*/me*^*4J*^ mice. Top: representative photograph of a 4-week-old female NOD *scid* +/*me*^*4J*^ (back) and NOD *scid*-*me*^*4J*^*/me*^*4J*^ (front) mouse, demonstrating the characteristic runted appearance and skin inflammation of homozygous mutants. Bottom: representative photograph of the hind foot of the same *me*^*4J*^*/me*^*4J*^ mouse, showing severe paw swelling and erythema. **(B)** Representative H&E-stained histological sections of paws from NOD *scid* +/+ (top) and NOD *scid*-*me*^*4J*^*/me*^*4J*^ (bottom) mice, demonstrating the dense suppurative inflammatory infiltrate in homozygous mutants. **(C)** Body weight of NOD *scid* +*/me*^*4J*^ and *scid*-*me*^*4J*^*/me*^*4J*^ mice at 3 weeks of age. n=7 (+/*me*^*4J*^) and n = 5 (*me*^*4J*^*/me*^*4J*^) *p < 0.05 by unpaired Student’s t-test. **(D)** Footpad inflammation score in NOD *scid* +*/me*^*4J*^ and *scid*-*me*^*4J*^*/me*^*4J*^ mice at 4 weeks of age. n=3 (+*/me*^*4J*^) and n=6 (*me*^*4J*^*/me*^*4J*^) **p < 0.01 by unpaired Student’s t-test. **(E)** Kaplan-Meier survival curves for NOD *scid me*^*4J*^*/me*^*4J*^ females (black; median survival 56.5 days; n=12) and males (red; median survival 53 days; n=19). No significant difference in survival was observed between sexes (log-rank test, p > 0.05), demonstrating that the lethal autoinflammatory disease affects both sexes with equivalent severity and kinetics. **(F)** Molecular characterization of the *me4J* mutation. Top left: schematic of the SHP-1 protein domain structure (N-SH2, C-SH2, and PTPase catalytic domain), with the location of the *me4J* mutation indicated within the catalytic domain. Bottom left: DNA sequence alignment showing the wild-type sequence (top) and the *me4J* mutant sequence (bottom), in which a single guanine (G) base insertion causes a frameshift leading to a premature stop codon (STOP), truncating the protein within the catalytic domain. Right: Sanger sequencing chromatograms from +/+, +*/me*^*4J*^, and *me*^*4J*^*/me*^*4J*^ mice confirming the insertion. Bottom: representative western blot showing total SHP-1 protein expression (anti-SHP-1, clone Y476) in BM lysates from +/+, +*/me*^*4J*^, and *me*^*4J*^*/me*^*4J*^ mice. GAPDH served as a loading control. SHP-1 protein is absent in *me*^*4J*^*/me*^*4J*^ mice and reduced in +*/me*^*4J*^ heterozygotes, confirming the loss-of-function nature of the mutation. **(G)** Splenomegaly in NOD *scid me*^*4J*^*/me*^*4J*^ mice. Top: representative photograph of spleens from +*/me*^*4J*^ and *me*^*4J*^*/me*^*4J*^ mice. Bottom: spleen weight expressed as a percentage of body weight. n = 4 (+*/me*^*4J*^) and n=7 (*me*^*4J*^*/me*^*4J*^). ****p < 0.0001 by unpaired Student’s t-test. **(H)** Representative H&E-stained histological sections of spleens from +*/me*^*4J*^ (left) and *me*^*4J*^*/me*^*4J*^ (right) mice, demonstrating disrupted splenic architecture and expanded red pulp in homozygous mutants. **(I)** Representative H&E-stained histological sections of kidneys from +*/me*^*4J*^ (left) and *me*^*4J*^*/me*^*4J*^ (right) mice, demonstrating glomerular and interstitial inflammatory infiltrates in homozygous mutants. **(J)** Serum concentrations of IL-6 and TNFα in 6-week-old +/+, +*/me*^*4J*^, and *me*^*4J*^*/me*^*4J*^ mice, measured by ELISA. Data are mean ± SEM from n=4 biological samples per group, each measured in duplicate. *p < 0.05, **p < 0.01 by unpaired Student’s t-test. **(K)** Quantification of neutrophil extracellular traps (NETs) in peripheral blood of NOD *scid* +*/me*^*4J*^ and *me*^*4J*^*/me*^*4J*^ mice at 4 weeks of age, assessed by SYTOX Orange staining of extracellular DNA. See Methods for details. n=8 (+*/me*^*4J*^) and n=7 (*me*^*4J*^*/me*^*4J*^). ***p < 0.001 by unpaired Student’s t-test. **(L)** Percentage of ApoTracker+ neutrophils in peripheral blood of NOD *scid* +*/me*^*4J*^ and *me*^*4J*^*/me*^*4J*^ mice at 6 weeks of age, reflecting the proportion of neutrophils undergoing apoptosis. n=6 per group. **p < 0.01 by unpaired Student’s t-test. **(M)** Schematic model summarizing the role of SHP-1 in regulating neutrophil activation. Loss of SHP-1 activity in *me*^*4J*^*/me*^*4J*^ neutrophils leads to increased pro-inflammatory cytokine (PIC) secretion and systemic inflammation.

Genetic mapping and Sanger sequencing identified a single guanine insertion in exon 7 of *Ptpn6* (CCGC→CCGGC) as the causative mutation. This insertion produces a frameshift beginning at Tyr278 that introduces a premature stop codon, truncating SHP-1 within its PTPase catalytic domain and resulting in loss of SHP-1 protein expression (**Figure 4F**). Consistent with loss of SHP-1–mediated restraint of cytokine signaling, homozygous mutants developed severe systemic inflammation, including massive splenomegaly with disrupted splenic architecture, widespread myeloid infiltration of the kidney, markedly elevated serum IL-6 and TNFα concentrations, and reduced body weight relative to heterozygous controls (**Figures 4G–4J**).

Circulating neutrophils from homozygous mutants displayed significantly elevated neutrophil extracellular trap (NET) formation and an elevated frequency of ApoTracker^+^ apoptotic cells compared with +/*me*^*4J*^ controls (**Figures 4K and 4L**), indicating that *me*^*4J/*^*me*^*4J*^ neutrophils are both hyperactivated and exhibit elevated apoptosis, consistent with activation-induced cell death, *in vivo*. Taken together, these data establish the *me4J* allele as a spontaneous SHP-1 loss-of-function mutation that causes severe, neutrophil-dominant autoinflammatory disease in the NOD *scid* background (**Figure 4M**). A companion study details the characterization of the *me4J* mutation on an immunocompetent NOD background, where it drives systemic autoimmunity (Arayan et al., submitted). Because SHP-1 acts as a phosphatase brake on multiple cytokine receptor signaling pathways—including those downstream of IL-1R and γc—this model provides an ideal system in which to test whether the IL-1/γc/MEK-ERK licensing axis identified *in vitro* contributes to pathological neutrophil activation and tissue damage *in vivo*.

### γc cytokine signaling is necessary and sufficient to drive autoinflammatory disease *in vivo*

To investigate whether γc cytokine signaling contributes to disease pathogenesis in the *me*^*4J/*^*me*^*4J*^ model, we first examined whether SHP-1–deficient neutrophils exhibit evidence of enhanced γc pathway engagement. Bulk RNA sequencing of flow-sorted bone marrow and peripheral blood neutrophils revealed that, relative to NOD scid +/+ controls (full transcriptomic analyses in Supplemental Figures 3 and 4), *me*^*4J/*^*me*^*4J*^ neutrophils coordinately upregulated multiple γc receptor subunit genes—including *Il2rb, Il15ra, Il7r*, and *Il21r*—in both compartments (**Figures 5A and 5B**; see **Figure 5C** for γc cytokines and receptor subunits). This coordinate transcriptional upregulation is consistent with increased sensitivity to γc cytokine signaling in the context of SHP-1 deficiency.

**Fig. 5.**
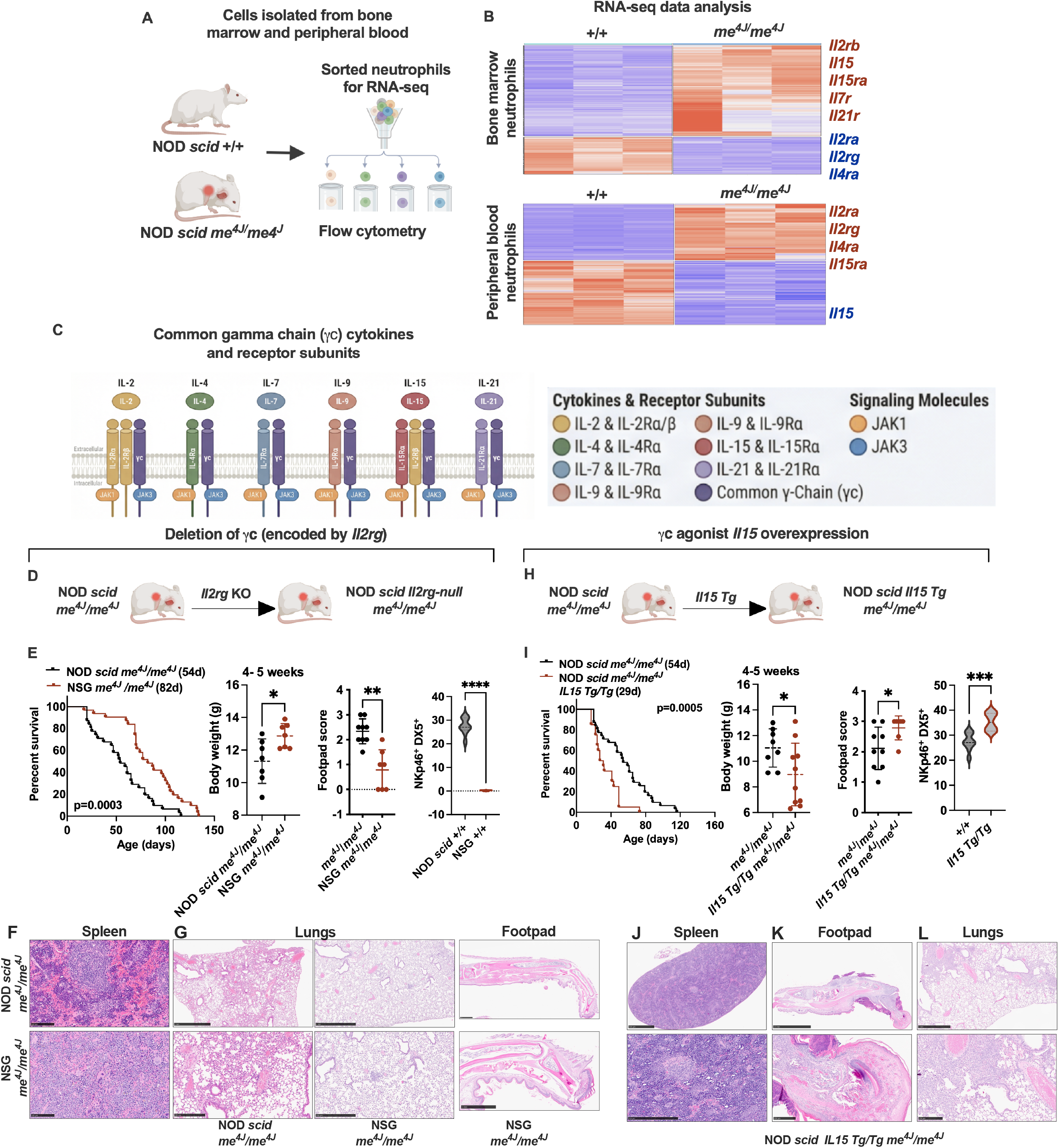
γc cytokine signaling is both necessary and sufficient to drive autoinflammatory disease in SHP-1-deficient mice. **(A)** Schematic of the experimental workflow for RNA-seq analysis. Bone marrow (BM) and peripheral blood (PB) neutrophils (Ly6G^+^CD11b^+^) were flow-sorted from NOD *scid*^*+/+*^ and NOD *scid*-*me*^*4J*^*/me*^*4J*^ mice for bulk RNA sequencing. **(B)** Heatmaps of differentially expressed genes from RNA-seq analysis of flow-sorted Ly6G+CD11b+ BM neutrophils (top) and PB neutrophils (bottom) from NOD scid +/+ and NOD *scid*-*me*^*4J*^*/me*^*4J*^ mice (n=3 per group). Heatmap colors represent row-normalized expression (z-score), with red indicating high and blue indicating low relative expression. Selected γc cytokine receptor subunit genes are annotated; gene names in red are significantly upregulated and those in blue are significantly downregulated in *me*^*4J*^*/me*^*4J*^ neutrophils (adjusted p < 0.05, |log2FC| > 1). Notably, transcripts encoding multiple γc receptor subunits—including *Il2rb, Il15, Il15ra*, and *Il7r*—are coordinately upregulated in *me*^*4J*^*/me*^*4J*^ neutrophils in both compartments, suggesting that loss of SHP-1 promotes a positive feedback mechanism that primes them for γc cytokine stimulation. **(C)** Schematic illustrating the receptor-JAK-STAT signaling architecture of the six common γ-chain (γc) cytokines: IL-2, IL-4, IL-7, IL-9, IL-15, and IL-21. Each cytokine signals through a unique receptor complex that incorporates the shared γc subunit (*IL2rg*), coupling to JAK1 and/or JAK3 to activate distinct STAT transcription factors. (**D-G**) Genetic ablation of γc signaling ameliorates autoinflammatory disease in NOD *scid me*^*4J*^*/me*^*4J*^ mice. Top left: schematic illustrating the genetic cross of NOD *scid*-*me*^*4J*^*/me*^*4J*^ mice with *Il2rg*-null mice to generate NSG-*me*^*4J*^*/me*^*4J*^ mice. Survival curve: Kaplan-Meier analysis of NOD *scid me*^*4J*^*/me*^*4J*^ (median survival 54 days; n=31 mice) and NSG-*me*^*4J*^*/me*^*4J*^ mice (median survival 82 days; n=31 mice); log-rank (Mantel-Cox) test, p=0.0003. At 4–5 weeks of age, NSG-*me*^*4J*^*/me*^*4J*^ mice exhibited significantly greater body weight (n=7 per group; *p < 0.05), reduced spleen weight as a percentage of body weight (n=7 vs. n=4; ***p < 0.001), and complete absence of footpad inflammation (footpad score = 0; n=9 vs. n=7; **p < 0.01). NSG +/+ mice lack NKp46^+^DX5^+^ cells (E; n=7–9 mice per group; ****p < 0.0001, unpaired t-test), confirming γc-dependent NK cell development. All other comparisons (*p < 0.05, **p < 0.01, ***p < 0.001, unpaired t-test) are versus NOD *scid*-*me*^*4J*^*/me*^*4J*^. All comparisons by unpaired Student’s t-test. Representative H&E-stained histological sections of spleen (F) and lungs and footpad (G; three panels, left: lungs NOD *scid*-*me*^*4J*^*/me*^*4J*^, middle: lungs NSG-*me*^*4J*^*/me*^*4J*^, right: footpad NSG-*me*^*4J*^*/me*^*4J*^) mice. Histological analysis revealed that disease amelioration in NSG-*me*^*4J*^*/me*^*4J*^ mice was associated with preserved splenic architecture (F), markedly reduced pulmonary inflammatory infiltrates (G, middle panel), and histologically normal footpad tissue (G, right panel), in stark contrast to the extensive neutrophilic infiltration observed in NOD *scid*-*me*^*4J*^*/me*^*4J*^ mice. (**H-L**) Constitutive overexpression of the γc agonist IL-15 is sufficient to accelerate disease. Top left: schematic illustrating the genetic cross of NOD *scid*-*me*^*4J*^*/me*^*4J*^ mice with constitutive *IL15* transgenic mice to generate NOD *scid*-*me*^*4J*^*/me*^*4J*^ *IL15* Tg/Tg mice. Survival curve: Kaplan-Meier analysis of NOD *scid*-*me*^*4J*^*/me*^*4J*^ (median survival 54 days; n=31) and NOD *scid*-*me*^*4J*^*/me*^*4J*^ *IL15* Tg/Tg mice (median survival 29 days; n=20), representing an approximately 46% reduction in median survival; log-rank (Mantel-Cox) test, p=0.0005. At 4–5 weeks of age, NOD *scid*-*me*^*4J*^*/me*^*4J*^ *IL15* Tg/Tg mice exhibited significantly reduced body weight (n=8 vs. n=10; *p < 0.05), increased spleen weight as a percentage of body weight (n=7 vs. n=6; *p < 0.05), and elevated footpad inflammation scores (n=9 vs. n=6; *p < 0.05) compared to NOD *scid*-*me*^*4J*^*/me*^*4J*^ controls. All comparisons by unpaired Student’s t-test. Additionally, there was a significant expansion of NKp46^+^DX5^+^ cells compared to IL-15 Tg controls (I; ***p<0.001, unpaired t-test). Histological examination revealed exacerbated pathology in the IL-15 Tg/Tg mice, including severe disruption of splenic architecture (J), ulcerative dermatitis with leukocytoclastic vasculitis in the footpad (K), and extensive pulmonary capillary congestion with alveolar hemorrhage (L). Data in panels E and I are presented as mean ± SEM. *p < 0.05, **p < 0.01, ***p < 0.001.

To determine whether γc signaling is required for disease, we crossed NOD *scid*-*me*^*4J/*^*me*^*4J*^ (SHP-1 deficient) mice onto the *Il2rg*-null NSG background, which lacks the shared γc receptor chain and is therefore globally deficient in γc cytokine signaling (**Figure 5D**). NSG-*me*^*4J/*^*me*^*4J*^ (γc and SHP-1 deficient) mice exhibited significant amelioration of disease compared with γc-sufficient NOD *scid*-*me*^*4J/*^*me*^*4J*^ littermates, including markedly prolonged survival (median 82 vs. 54 days; p = 0.0003), improved body weight, reduced footpad inflammation scores, and substantially attenuated splenic pathology and pulmonary inflammatory infiltrates (**Figures 5E–G**). Histological examination of footpad tissue confirmed the absence of inflammatory pathology in NSG-*me*^*4J*^*/me*^*4J*^ mice, which displayed histologically normal footpad architecture with no evidence of the ulcerative dermatitis or leukocytic infiltration seen in NOD *scid-me*^*4J*^*/me*^*4J*^ controls (**Figure 5G**). As expected from the role of IL-15–γc signaling in NK cell development(*16*), NSG +/+ mice lacked NKp46^+^ DX5^+^ NK cells (**Figure 5E**). Although this precludes attribution of rescue exclusively to neutrophil-intrinsic γc signaling, the data establish that γc receptor engagement is necessary for full disease expression.

To ask whether augmented γc signaling is sufficient to accelerate disease, we crossed NOD *scid me*^*4J/*^*me*^*4J*^ mice to a transgenic line constitutively overexpressing IL-15 (**Figure 5H**). IL-15 Tg/Tg mice on a wild-type background remain overtly healthy; however, constitutive IL-15 signaling in the SHP-1–deficient context dramatically accelerated disease progression. NOD *scid* IL-15 Tg/Tg *me*^*4J/*^*me*^*4J*^ mice showed an approximately 46% reduction in median survival relative to non-transgenic *me*^*4J/*^*me*^*4J*^ littermates (29 vs. 54 days; p = 0.0005), accompanied by greater weight loss and more severe footpad inflammation at 4–5 weeks of age (**Figure 5I**). This accelerated disease was associated with marked NK cell expansion (**Figure 5I**), and histological analysis revealed severe systemic pathology including disrupted splenic architecture (**Figure 5J**), ulcerative dermatitis with leukocytoclastic vasculitis of the footpad (**Figure 5K**), and extensive pulmonary capillary congestion with alveolar hemorrhage (**Figure 5L**).

Together, these complementary loss- and gain-of-function genetic experiments demonstrate that γc cytokine signaling is both necessary and sufficient to modulate autoinflammatory disease severity in SHP-1– deficient mice, and that the magnitude of γc pathway activity scales directly with disease outcome. These findings extend our *in vitro* observations to an *in vivo* setting and implicate γc-dependent licensing of the IL-1/MEK-ERK axis as a pathologically relevant mechanism of neutrophil-driven inflammation.

### γc signaling is required for terminal neutrophil activation and inflammatory effector function

To define the cellular mechanism underlying the *in vivo* protection observed in γc-deficient mice, we first assessed the activation state of neutrophils from NOD *scid*-*me*^*4J/*^*me*^*4J*^ and NSG-*me*^*4J/*^*me*^*4J*^ animals. While bone marrow neutrophils from both genotypes were predominantly in a resting state (**Figures 6A and 6B**), a striking difference emerged at the site of inflammation. Neutrophils infiltrating the footpads of NOD *scid*-*me*^*4J/*^*me*^*4J*^ mice had largely transitioned to an activated phenotype (gating strategy in Supplemental Figure 5), whereas neutrophils from NSG-*me*^*4J/*^*me*^*4J*^ mice failed to undergo this transition and remained predominantly in an intermediate state, as illustrated by representative flow cytometry plots (**Figure 6C**) and corresponding quantification (**Figure 6D**). These results suggest that an intact γc signaling axis is required for neutrophil terminal activation *in vivo*. To establish that this requirement is cell-intrinsic and directly attributable to γc receptor signaling rather than indirect effects of the inflammatory microenvironment, we tested whether γc-deficient neutrophils fail to activate in response to defined cytokine stimulation *in vitro*. Co-stimulation of NOD scid +/+ BM neutrophils with IL-1α and IL-15 drove a dose- and time-dependent increase in the Activated population and a corresponding decrease in Resting neutrophils; this response was completely abolished in NSG +/+ BM neutrophils lacking the γc receptor chain (**Figures 6E–6G**), indicating that the γc receptor is cell-intrinsically required for IL-1α+IL-15-driven neutrophil activation independent of the *in vivo* inflammatory context.

**Fig. 6.**
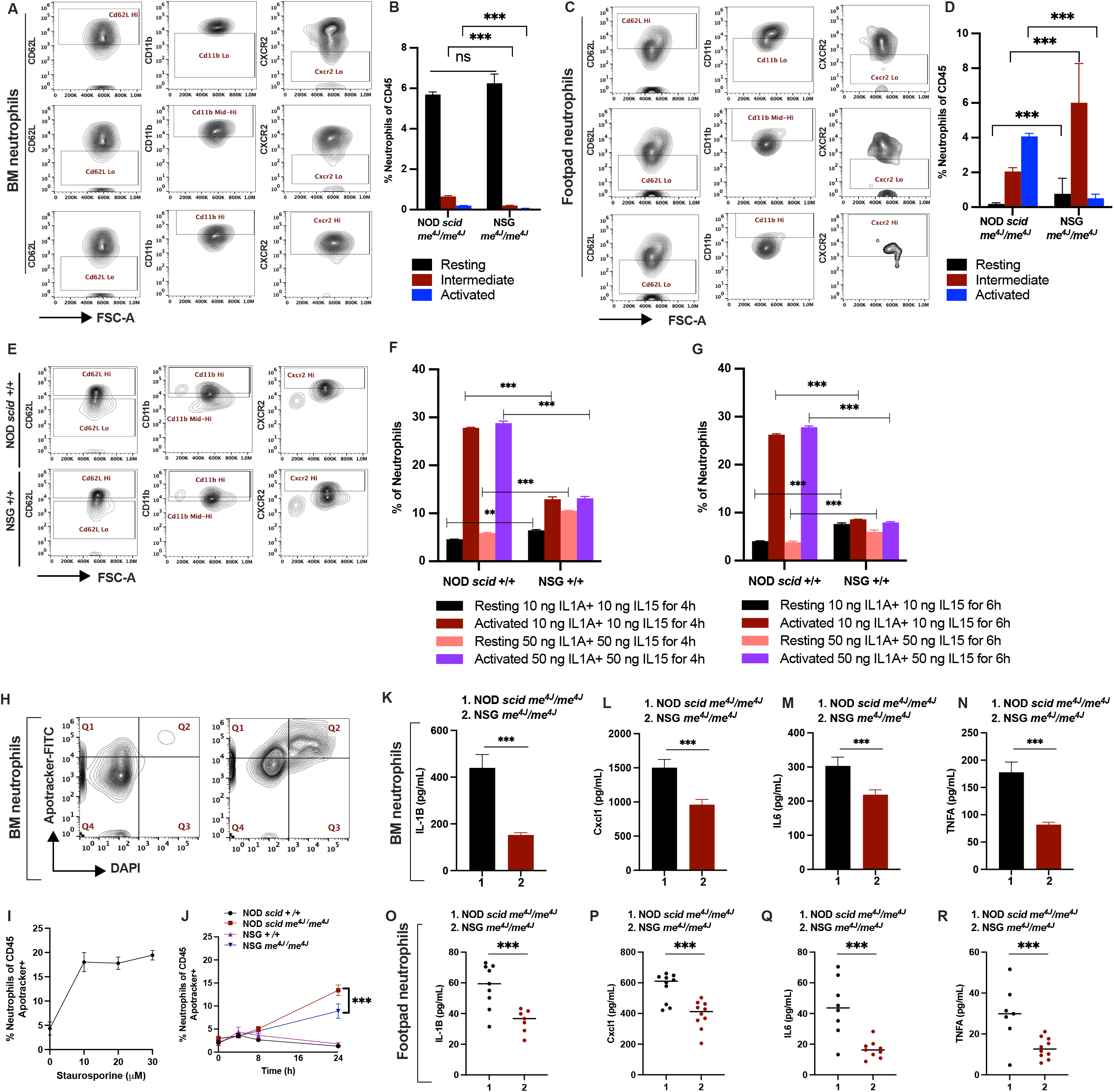
γc signaling is required for the full inflammatory activation, cytokine secretory capacity, and survival of neutrophils at the site of disease. **(A–D)** Neutrophil activation state phenotyping. Bone marrow (BM) neutrophils **(A, B)** and footpad neutrophils **(C, D)** from NOD *scid-me*^*4J*^*/me*^*4J*^ and NSG-*me*^*4J*^*/me*^*4J*^ mice were classified into three activation states by flow cytometry: Resting (CD62L^hi^ CD11b^lo^ CXCR2^lo^), Intermediate (CD62L^lo^ CD11b^mid-high^ CXCR2^lo^), and Activated (CD62L^lo^ CD11b^hi^ CXCR2^hi^). Representative flow cytometry plots are shown for BM **(A)** and footpad **(C)**. These gates operationally define activation states along a continuous spectrum of CD62L downregulation and CD11b/CXCR2 upregulation; the boundaries reflect empirically distinct surface marker expression patterns rather than discrete biological entities. In the BM **(B)**, both genotypes harbored predominantly Resting neutrophils, with a modest but significant reduction in the Intermediate and Activated populations in NSG-*me*^*4J*^*/me*^*4J*^ mice. In the footpad **(D)**, NOD *scid-me*^*4J*^*/me*^*4J*^ neutrophils were predominantly in the Activated state, whereas NSG-*me*^*4J*^*/me*^*4J*^ footpad neutrophils were predominantly Intermediate, demonstrating that γc signaling is required for neutrophils to transition to the fully activated, tissue-recruited phenotype at the site of inflammation (n=3 per group; ***p < 0.001, multiple unpaired t-tests). **(E–G)** γc receptor signaling is required for IL-1α+IL-15-driven neutrophil activation *in vitro*. BM neutrophils from NOD *scid* +/+ and NSG +/+ mice were co-stimulated with IL-1α + IL-15 at two doses (10 ng/mL or 50 ng/mL each) for 4 h **(E, F)** or 6 h **(G)**, and neutrophil activation state was assessed by flow cytometry using CD62L, CD11b, and CXCR2 surface markers (gating as in A–D). **(E)** Representative flow cytometry plots for the 4 h timepoint, shown for NOD *scid* +/+ (top) and NSG +/+ (bottom). **(F, G)** Quantification of the percentage of Resting and Activated neutrophils at 4 h (F) and 6 h (G). In NOD *scid* +/+ neutrophils, co-stimulation significantly increased the proportion of Activated neutrophils and decreased the Resting population in a dose-dependent manner at both timepoints; this response was abolished in NSG +/+ neutrophils, which lack the shared γc receptor chain, demonstrating that γc receptor signaling is cell-intrinsically required for IL-1α+IL-15-driven neutrophil activation *in vitro*. Data are mean ± SEM; n=3 biological replicates per group. **p < 0.01, ***p < 0.001 by unpaired t-test. **(H)** Representative flow cytometry plots of ApoTracker/DAPI staining in BM neutrophils, used to assess apoptosis. **(I)** Validation of the ApoTracker apoptosis assay. BM neutrophils from NOD *scid* +/+ mice were treated with increasing concentrations of staurosporine (0–30 μM), a broad kinase inhibitor used as a positive control for apoptosis induction. The percentage of ApoTracker+ neutrophils (gated as Ly6G+CD11b+) increased in a dose-dependent manner, confirming assay sensitivity (n=3, each measured in duplicate). **(J)** Spontaneous apoptosis time course. BM neutrophils from NOD *scid* +/+, NOD *scid*-*me*^*4J*^*/me*^*4J*^, NSG +/+, and NSG-*me*^*4J*^*/me*^*4J*^ mice were cultured *ex vivo* without stimulation for up to 24 h, and the percentage of ApoTracker+ neutrophils was measured at indicated time points (n=3 per group). At 24 h, NOD *scid*-*me*^*4J*^*/me*^*4J*^ neutrophils exhibited significantly greater spontaneous apoptosis than NSG-*me*^*4J*^*/me*^*4J*^ neutrophils (***p < 0.001, unpaired t-test), consistent with a more terminally activated state driven by intact γc signaling. **(K–N)** Bone marrow (BM) neutrophils were flow-sorted (Ly6G+CD11b+CD45+) from 8-week-old NOD *scid*-*me*^*4J*^*/me*^*4J*^ (group 1) and NSG-*me*^*4J*^*/me*^*4J*^ (group 2) mice and stimulated *ex vivo* with low-dose LPS (10 ng/mL) for 8 h. Supernatant concentrations of IL-1β **(K)**, CXCL1 **(L)**, IL-6 **(M)**, and TNFα **(N)** were significantly reduced in NSG-*me*^*4J*^*/me*^*4J*^ BM neutrophils compared to NOD *scid*-*me*^*4J*^*/me*^*4J*^ controls (n=3 per group, each measured in duplicate by ELISA; ***p < 0.001, unpaired t-test). **(O–R)** Neutrophils were flow-sorted from the footpads of 8-week-old NOD *scid*-*me*^*4J*^*/me*^*4J*^ (group 1) and NSG-*me*^*4J*^*/me*^*4J*^ (group 2) mice and stimulated *ex vivo* with LPS (10 ng/mL) for 8 h. Footpad neutrophils, which represent tissue-infiltrating cells that have undergone partial activation *in vivo*, exhibited lower baseline secretory capacity than BM neutrophils; nevertheless, concentrations of IL-1β **(O)**, CXCL1 **(P)**, IL-6 **(Q)**, and TNFα **(R)** were significantly reduced in NSG-*me*^*4J*^*/me*^*4J*^ footpad neutrophils compared to NOD *scid*-*me*^*4J*^*/me*^*4J*^ controls (n=3 per group, each measured in duplicate by ELISA; ***p < 0.001, unpaired t-test).

To evaluate whether this failure to reach terminal activation impacted neutrophil survival, we first validated an ApoTracker apoptosis assay by stimulating bone marrow neutrophils with staurosporine as a positive control, yielding representative flow cytometry plots (**Figure 6H**) and dose-dependent quantification (**Figure 6I**). Using this assay, we then measured spontaneous apoptosis in bone marrow-derived neutrophils from the four indicated strains. Consistent with a failure to reach terminal activation, NSG-*me*^*4J*^*/me*^*4J*^ neutrophils exhibited significantly reduced spontaneous apoptosis during *ex vivo* culture compared to their counterparts from NOD *scid*-*me*^*4J*^*/me*^*4J*^ mice (**Figure 6J**). This is mechanistically consistent with a model where the synergistic signaling required for terminal activation also primes cells for activation-induced cell death; by failing to receive this signal *in vivo*, NSG neutrophils do not engage this pro-apoptotic program.

This defect in terminal activation was directly associated with a profound impairment of inflammatory effector function. Upon *ex vivo* stimulation, neutrophils sorted from the bone marrow of NSG-*me*^*4J*^*/me*^*4J*^ mice exhibited a markedly reduced capacity to secrete inflammatory cytokines, including IL-1β, CXCL1, IL-6, and TNFα, compared to their γc-sufficient counterparts (**Figures 6K–6N**). This functional impairment was even more pronounced in neutrophils isolated directly from sites of inflammation (**Figures 6O–6R**). Together, these results demonstrate that γc signaling is required for neutrophils to undergo full inflammatory activation and that, in its absence, neutrophils fail to transition into a terminally activated, cytokine-secreting state, providing a cellular explanation for the protection from autoinflammatory pathology observed in γc-deficient mice. This impairment encompassed all four cytokines measured—IL-1β, CXCL1, IL-6, and TNFα—in both bone marrow and footpad-infiltrating neutrophils, confirming that the effector function deficit is not restricted to a single mediator and is recapitulated at the site of inflammation.

### Individual γc cytokine pathways are dispensable in this autoinflammatory model

Our model predicts that multiple γc cytokines can sustain this licensing program. To test this hypothesis *in vivo*, we systematically crossed NOD *scid*-*me*^*4J*^*/me*^*4J*^ mice to strains lacking individual γc cytokine ligands or receptors. In stark contrast to the profound protection conferred by ablation of the shared γc receptor (**Figure 5D**), deletion of the individual γc cytokine pathways tested here had no detectable effect on disease progression. Genetic ablation of the receptors or ligands for IL-2/IL-15 (via *Il2rb*), IL-4, IL-7 (*Il7r*), IL-9 (*Il9r*), IL-15, or IL-21 (*Il21r*) failed to alter disease kinetics, survival, body weight, or inflammatory pathology in NOD *scid*-*me*^*4J*^*/me*^*4J*^ mice (**Figures 7A–F**). Notably, deletion of *Il2rb* or *Il15* resulted in complete loss of NK cells, confirming effective disruption of IL-15–dependent NK cell development (**Figures 7A and 7E**). Despite this profound NK cell depletion, disease progression remained unchanged. These findings demonstrate that NK cells are not required effectors in this model of autoinflammatory disease. Together, these results show that the tested γc cytokine pathways are functionally buffered *in vivo* in this model. While elimination of the shared γc receptor markedly attenuates disease, removal of any single tested γc cytokine pathway is insufficient to disrupt the inflammatory program.

**Fig. 7.**
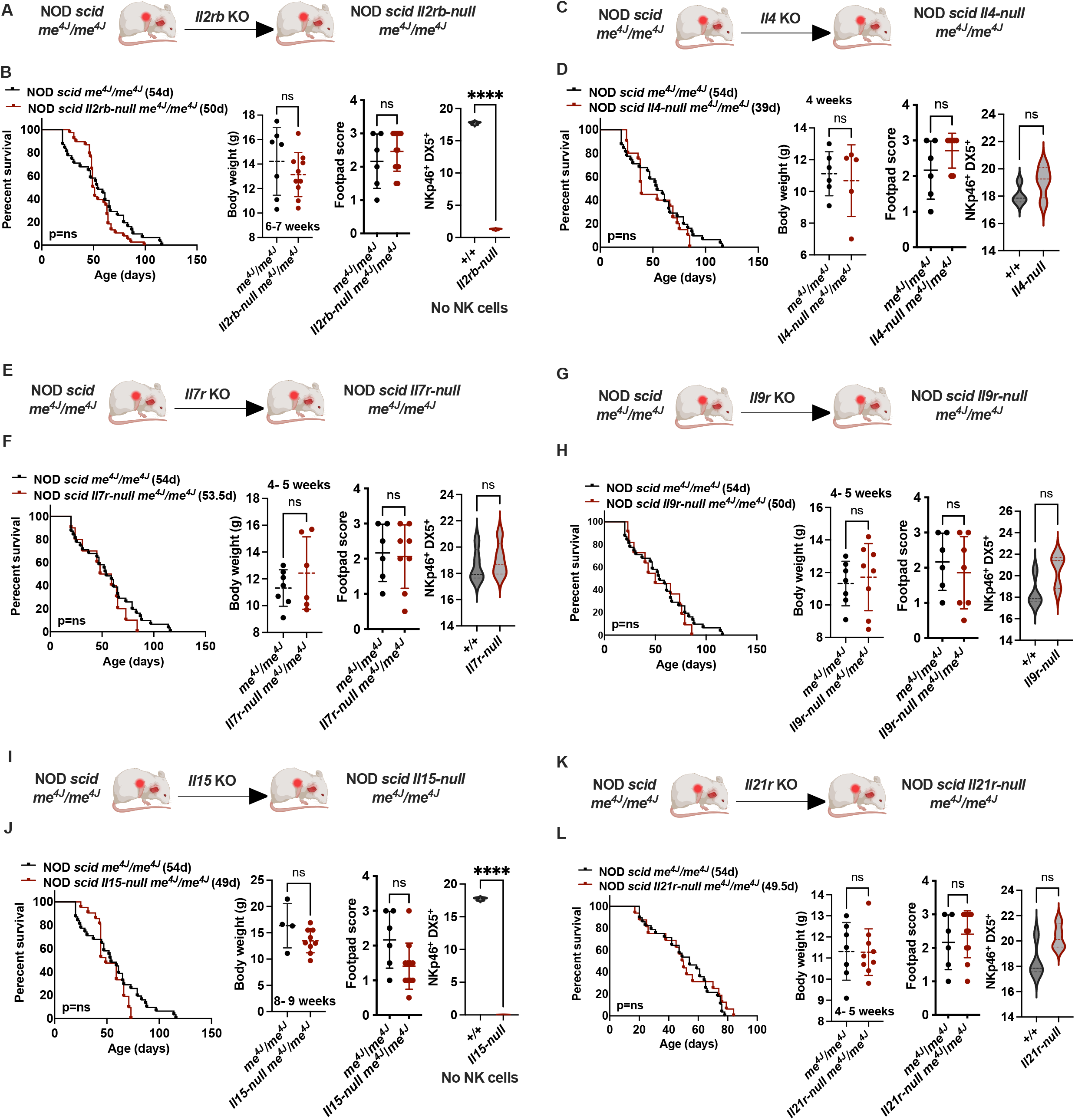
Individual γc cytokines are redundant for the development of autoinflammatory disease. **(A)** NOD *scid*-*me*^*4J*^*/me*^*4J*^ mice were crossed to *Il2rb*-null mice, which lack IL-2Rβ, a shared receptor subunit for IL-2 and IL-15. Kaplan-Meier survival analysis (n=31 vs. 38 mice), body weight, and footpad score at 6–7 weeks were not significantly different from NOD *scid*-*me*^*4J*^*/me*^*4J*^. Peripheral blood NKp46+DX5+ NK cells were absent in *Il2rb*-null mice compared to wild-type controls (n=4 per group, ****p < 0.0001), confirming successful NK cell depletion; the lack of disease rescue demonstrates that NK cells are not required effectors in this model. **(B)** NOD *scid*-*me*^*4J*^*/me*^*4J*^ mice were crossed to *Il4*-null mice. Survival (n=31 vs. 20 mice), body weight, and footpad score at 4 weeks were not significantly different from controls. NKp46+DX5+NK cells were present at normal levels in *Il4*-null mice (n=4 mice per group, ns). **(C)** NOD *scid*-*me*^*4J*^*/me*^*4J*^ mice were crossed to *Il7r*-null mice. Survival (n=31 vs. 10 mice), body weight, and footpad score at 4–5 weeks were not significantly different from controls. NKp46+DX5+NK cells were present at normal levels in *Il7r*-null mice (n=4 per group, ns). **(D)** NOD *scid*-*me*^*4J*^*/me*^*4J*^ mice were crossed to *Il9r*-null mice. Survival (n=31 vs. 11 mice), body weight, and footpad score at 4–5 weeks were not significantly different from controls. NKp46+DX5+NK cells were present at normal levels in *Il9r*-null mice (n=4 per group, ns). **(E)** NOD *scid*-*me*^*4J*^*/me*^*4J*^ mice were crossed to *Il15*-null mice. Survival (n=31 vs. 21 mice), body weight, and footpad score at 8–9 weeks were not significantly different from controls. Peripheral blood NKp46+DX5+NK cells were absent in *Il15*-null mice compared to wild-type controls (n=4 per group, ****p < 0.0001), confirming NK cell depletion; the lack of disease rescue further demonstrates that NK cells are dispensable for autoinflammatory disease progression. **(F)** NOD *scid*-*me*^*4J*^*/me*^*4J*^ mice were crossed to *Il21r*-null mice. Survival (n=31 vs. 13 mice), body weight, and footpad score at 4–5 weeks were not significantly different from controls. NKp46+DX5+NK cells were present at normal levels in *Il21r*-null mice (n=4 per group, ns). Data are presented as mean ± SEM. Survival curves were compared using the Log-rank (Mantel-Cox) test. All other comparisons were made using an unpaired t-test. ns, not significant; *, p < 0.05; ****, p < 0.0001.

## DISCUSSION

Prior genetic dissection of *Ptpn6*-dependent autoinflammation has firmly established IL-1α as the central cytokine driving neutrophilic disease, with a well-defined cascade of downstream effectors—including SYK, MyD88, RIPK1, TAK1, CARD9, and NFκB—elucidated through systematic knockout approaches(*8, 14, 17, 18*). Despite these advances, the upstream mechanism that licenses IL-1α to initiate and amplify this effector program at sites of inflammation remained unresolved. Here, we identify the common γ-chain (γc) cytokine family as this critical licensing tier. Multiple γc cytokines can provide this signal interchangeably, while the downstream response prominently involves MEK/ERK signaling. This organization helps explain how inflammatory licensing remains robust across contexts despite variability in the specific upstream γc cytokines present.

The overlap observed among γc cytokines aligns with the fundamental biology of the common γ-chain receptor system. The γc receptor (encoded by *IL2RG*) functions as a shared signaling subunit for IL-2, IL-4, IL-7, IL-9, IL-15, and IL-21, enabling diverse ligands to activate overlapping downstream pathways predominantly through JAK–STAT modules(*6, 19*). This receptor-sharing architecture has long been recognized as a source of robustness within lymphocyte cytokine networks. The present data extend this principle to innate immunity by showing that a similar overlap preserves IL-1–driven inflammatory permissiveness even when individual γc cytokine inputs are absent. The common γc receptor itself emerges as the shared indispensable component, with its genetic ablation—rather than loss of any single ligand—being necessary to disrupt licensing *in vivo*.

Although γc cytokines are classically associated with lymphocyte biology, neutrophils express functional γc receptor subunits and respond directly to γc cytokine stimulation. IL-15 enhances neutrophil phagocytic activity, delays apoptosis, and activates ERK1/2 through a Syk-dependent mechanism in primary human neutrophils(*20-22*). The present data extend this biology to a synergistic co-stimulation context: IL-1α and γc signals together produce MEK/ERK activation exceeding the additive effects of either input alone. Mechanistically, the substantial suppression of cytokine output by MEK inhibition via trametinib, contrasted with only partial attenuation by JAK3/STAT5 inhibition, indicates that MEK/ERK functions downstream of—or parallel to—STAT5 signaling rather than as a simple effector of either pathway independently(*23*). This pharmacological dissection distinguishes the MEK/ERK-dependent response described here from the p38 MAPK pathway previously characterized in Ptpn6ΔPMN neutrophils, which promotes TNF and IL-1α/β production through a cell-intrinsic, RIPK1-independent mechanism(*8, 24*). Together, these findings suggest that *Ptpn6*-dependent neutrophil inflammation is governed by two distinct MAPK tiers: a p38 arm controlling basal inflammatory tone intrinsically, and a MEK/ERK arm that amplifies inflammatory output upon γc co-stimulation within the tissue microenvironment. The NOD *scid*-*me*^*4J*^*/me*^*4J*^ model—harboring a frameshift allele that produces complete SHP-1 loss in an inflammatory-sustaining background—reveals this MEK/ERK-dependent component more completely than partial-loss models.

A formal limitation of our genetic approach warrants acknowledgment: the rescue conferred by *Il2rg* ablation reflects global loss of the γc receptor chain and does not, in principle, exclude contributions from non-canonical γc biology beyond the six classically defined cytokine receptor complexes. The γc chain is a structurally dynamic protein whose receptor-sharing function extends beyond cytokine-induced ectodomain heterodimerization. Recent structural work has demonstrated that the γc transmembrane domain plays an active, ligand-independent role in receptor assembly, engaging the transmembrane domains of diverse interleukin receptors through a conserved knob-into-hole mechanism within the lipid bilayer itself—a mode of interaction capable in principle of nucleating receptor assemblies not captured by individual cytokine knockouts (*25*). Additionally, alternative splicing of the *Il2rg* pre-mRNA generates a secreted soluble γc isoform (sγc) that homodimerizes and competitively associates with IL-2Rβ at the cell surface, modulating signaling output in a JAK3- and ligand-independent manner, and the intracellular domain of γc harbors four tyrosine residues of incompletely characterized function, leaving open the possibility of outputs not fully mediated through the canonical JAK3-STAT axis(*19*). However, several converging lines of evidence argue strongly that canonical γc cytokine signaling, rather than a non-canonical γc function, accounts for our findings. Most compellingly, constitutive overexpression of IL-15 alone—a single canonical γc ligand operating exclusively through its defined receptor complex—is sufficient to dramatically accelerate disease in *me*^*4J*^*/me*^*4J*^ mice, reducing median survival by approximately 46% and exacerbating histopathological severity across multiple tissues. This gain-of-function result demonstrates that quantitative augmentation of conventional γc cytokine input directly scales disease output, a relationship that is not predicted by non-canonical γc mechanisms but is precisely what the canonical licensing model requires. Complementing this, the complete panel of individual γc cytokine receptor knockouts—encompassing all licensing-competent family members—fails to phenocopy the protection conferred by global *Il2rg* ablation, a result fully explained by functional redundancy among canonical inputs. Furthermore, the *in vitro* reconstitution experiments show that γc-deficient NSG neutrophils fail to respond to defined IL-1α plus IL-15 co-stimulation in a purified cytokine system, directly localizing the γc chain requirement to its role in canonical receptor signaling rather than to indirect or structural effects. Taken together, the bidirectional dose-dependence of disease on γc cytokine ligand availability—protection upon receptor ablation, acceleration upon ligand excess—constitutes strong functional evidence that the γc licensing circuit operates through canonical cytokine-receptor engagement. Definitive exclusion of non-canonical contributions would require neutrophil-specific conditional re-expression of signaling-competent versus signaling-dead γc alleles, which remains an important avenue for future investigation.

The *in vivo* genetic experiments are consistent with γc signaling operating through a neutrophil-intrinsic mechanism, though formal cell-autonomous proof would require conditional neutrophil-specific deletion of Il2rg. A transition to a hyperactivated, CD62L^lo^ state in peripheral tissues is a core feature of the *me4J* allele and has likewise been observed in the spleen of immunocompetent NOD *me*^*4J*^*/me*^*4J*^ mice (Arayan et al., submitted). To test whether this phenotype reflected cell-autonomous γc signaling or indirect effects mediated by γc-dependent lymphoid populations, individual cytokine pathway knockouts were evaluated. Deletion of *Il2rb* and *Il15*—genes essential for NK cell development and homeostasis—failed to protect *me*^*4J*^*/me*^*4J*^ mice despite confirmed NK cell depletion, demonstrating that NK cells are dispensable effectors in this model. Notably, the original SHP-1-deficient motheaten mice also exhibited virtually absent NK cell activity, yet still developed severe autoinflammation (*26*). These results rule out NK cells as required effectors, but because the NSG cross ablates γc signaling globally, the data are consistent with—but do not formally prove—a neutrophil-intrinsic mechanism. Consistent with this, flow cytometric analysis of CD62L, CD11b, and CXCR2 expression demonstrated that NSG-*me*^*4J*^*/me*^*4J*^ footpad neutrophils remained predominantly in an intermediate, rather than activated, state, directly localizing the γc requirement to the step of terminal activation at the inflammatory site rather than to neutrophil recruitment per se.

The sensitivity of this licensing circuit is mechanistically grounded in SHP-1’s role as a direct negative regulator of the γc receptor signaling complex. Upon γc cytokine stimulation, SHP-1 is recruited to the assembled receptor complex—comprising the γc chain, IL-2Rβ, and their associated kinases—where it directly dephosphorylates JAK1 (constitutively associated with IL-2Rβ) and JAK3 (recruited to γc upon ligand binding), as well as IL-2Rβ phospho-tyrosine residues themselves(*27*). Importantly, this dephosphorylation is optimal only when the full receptor complex is assembled and is dependent on SHP-1’s phosphatase activity, as demonstrated by the failure of a phosphatase-dead SHP-1 mutant to suppress IL-2 receptor phosphorylation (*27*). SHP-1 thus functions as a feedback brake specifically calibrated to the assembled, active γc signaling platform rather than to individual cytokine pathways in isolation(*5*). In the absence of SHP-1, JAK3 activation at the γc complex becomes sustained, receptor phospho-tyrosines remain occupied, and the downstream MEK/ERK cascade is no longer subject to the self-limiting kinetics that characterize physiological γc cytokine signaling(*5, 8*). This amplified and sustained γc signal, converging with IL-1α input on the MEK/ERK node, drives the pathological inflammatory output observed in *me*^*4J/*^*me*^*4J*^ mice. Of note, MEK and ERK can be activated independently of one another in human neutrophils under certain stimulatory conditions (*28*), underscoring that the substantial suppression of cytokine output by trametinib observed here is consistent with MEK-dependent ERK activation as a dominant route in the γc/IL-1α co-stimulation context. This framework is complementary to—rather than in competition with—the p38 and caspase-8–dependent mechanisms previously shown to govern IL-1α/β processing and release from *Ptpn6*-deficient neutrophils(*8*): whereas those mechanisms operate at the level of IL-1 production and secretion, the γc/MEK-ERK axis governs the amplitude of IL-1–driven effector activation downstream of IL-1 receptor engagement.

The licensing framework identified here complements prior observations by defining the positive regulatory signals that convert intrinsically hyperactive SHP-1–deficient neutrophils into fully pathogenic effectors *in vivo*. Mazgaeen et al. demonstrated that transfer of Ptpn6-deficient bone marrow into CD47-deficient recipients induces fatal IL-1–dependent systemic inflammation, establishing that intrinsically hyperactive SHP-1–deficient neutrophils are normally restrained by inhibitory signals from the host tissue environment(*14*). The present study identifies the complementary positive signals required for full inflammatory activation: γc cytokines present in the tissue microenvironment that amplify IL-1–driven output through MEK/ERK. The spontaneous *me4J* model further extends these observations by demonstrating that γc signaling is not merely permissive but actively disease-modifying—IL-15 transgene expression accelerates lethal disease, while genetic γc ablation prolongs survival—confirming that this pathway has substantial pathological consequence *in vivo*.

These findings carry direct implications for the treatment of IL-1–driven inflammatory diseases. The genetic data indicate that blockade of any single γc cytokine will likely have limited therapeutic efficacy, because compensatory signaling through the shared γc receptor can maintain the licensing circuit. This prediction aligns with the incomplete efficacy of cytokine-targeted therapies in complex inflammatory diseases such as sJIA, where IL-1 blockade reduces systemic inflammation but does not consistently normalize neutrophil biology(*2*). The more tractable intervention points may be the shared γc receptor and downstream MEK/ERK signaling, with direct *in vivo* support for the former and strong mechanistic support for the latter. The therapeutic case for MEK/ERK inhibition is further strengthened by evidence that this pathway integrates multiple inflammatory inputs in human neutrophils. Phospho-proteomic profiling of primary human neutrophils(*15*) demonstrates that LPS, which signals through the same MyD88/IRAK/TRAF6 axis as IL-1R, induces only modest ERK phosphorylation — mirroring the weak pERK response to IL-1α alone in our mouse model — further supporting that TLR4/IL-1R-type stimulation requires a co-stimulatory input to achieve full MEK/ERK activation (Figure 3K)(*8*). This indicates that MEK/ERK functions as a broader integration hub for inflammatory inputs in human neutrophils. In complex inflammatory diseases where multiple cytokine and innate immune pathways are simultaneously active, this downstream position may offer advantages over blockade of individual upstream cytokines. The translational relevance of this circuit is supported by our re-analysis of public transcriptomic datasets(*3*) from sJIA neutrophils and hidradenitis suppurativa skin lesions (**Figure 1**), which demonstrated coordinated upregulation of IL-1, γc, and MEK/ERK pathway components in human autoinflammatory disease (*9, 10*). Separately, the primary literature in these diseases has reported broad inflammatory and JAK/STAT-associated pathway activation in disease-relevant samples(*11, 12*). Together, these data provide a mechanistic rationale for testing downstream signaling hubs, rather than individual upstream cytokines, in settings where cytokine network redundancy limits the efficacy of cytokine-specific approaches. Critically, the cellular basis for this principle is established at the level of neutrophil activation state: γc-deficient neutrophils reach the inflamed tissue but fail to transition to the CD62Llo CD11bhi activated phenotype, consistent with a model in which downstream MEK/ERK-dependent signaling governs terminal inflammatory activation more than initial recruitment.

The same *me4J* allele, characterized on the immunocompetent NOD background in a companion study (Arayan et al., submitted), produces a distinct but complementary disease trajectory: when SHP-1-deficient neutrophils operate within a complete immune system, the same hyperactivated signaling state that drives neutrophil-intrinsic IL-1/MEK-ERK inflammation also drives systemic autoimmunity (Arayan et al., submitted). Together, the two studies define complementary windows into the same underlying biology: neutrophil-intrinsic signaling governs the amplitude and character of the innate inflammatory response, while the genetic immune context—specifically the presence or absence of B cell competence—determines whether that response escalates to systemic autoimmunity.

In summary, our findings reveal that IL-1–driven neutrophil inflammation depends on a permissive γc cytokine network that amplifies downstream MEK/ERK signaling. Because no single γc cytokine is essential, blockade of individual family members is unlikely to fully disrupt this program. By contrast, shared γc signaling *in vivo* and downstream MEK/ERK activation provide a mechanistic framework for understanding why single-cytokine interventions can underperform in complex inflammatory diseases and highlight shared signaling components as more promising therapeutic targets.

## MATERIALS AND METHODS

### Mice

All mice were housed under specific pathogen-free conditions at the Jackson Laboratory. Animal experiments were conducted under protocols approved by the Institutional Animal Care and Use Committee (IACUC protocols # 22022, 99099). Both male and female mice were used. The *Ptpn6 me*^*4J*^ allele—a frameshift mutation arising from a spontaneous guanine insertion in exon 7 **of** *Ptpn6*—was identified during routine colony maintenance of NOD *scid* mice (JAX: 001303). NOD-*me*^*4J*^ mice were maintained as heterozygotes on the NOD/ShiLtJ background and intercrossed to generate homozygous *Ptpn6 me*^*4J*^/*me*^*4J*^ (NOD-*me*^*4J*^) offspring.

### SNP identification

The *me*^*4J*^ allele was identified by Sanger sequencing of the *Ptpn6* locus. Genomic DNA was extracted from tail clips using standard proteinase K digestion. The *Ptpn6* coding sequence was amplified by PCR using Advantage-GC2 PCR Kit (Clontech/Takara, Cat# 639119) with the primers listed in the Key Resources Table. PCR products were purified with QIAquick Gel Extraction Kit (Qiagen, Cat# 28704) and treated with ExoSAP-IT (Thermo Fisher, Cat# 78200). Sanger sequencing was performed on an ABI 3730xl DNA Analyzer using BigDye Terminator v3.1 (Applied Biosystems, Cat# 4337455). Sequences were assembled and analyzed using Sequencher 4.9.

### Western blot

Spleens were collected from three mice per genotype (+/+, +/*me*^*4J*^, and *me*^*4J*^*/me*^*4J*^) at 6 weeks of age. Total protein lysates were prepared in Tris-buffered saline containing 1% Igepal and protease inhibitors on ice. Equal amounts of protein were resolved on 4–12% Tris-glycine precast gels and transferred using the iBLOT system (Invitrogen). Membranes were blocked in 5% non-fat dry milk in TBS containing 0.05% Tween-20 and incubated overnight at 4°C with rabbit anti-mouse PTPN6 (Abcam, Cat # ab32559; 1:500) or rabbit anti-mouse GAPDH (Cell Signaling Technology, Cat # 2118; 1:1000). HRP-conjugated secondary antibody (Cell Signaling Technology, Cat # 7074) was applied for 1 hour at room temperature. Signal was developed with West Pico PLUS Chemiluminescent Substrate (Thermo Fisher, Cat # 34577) and recorded using a CCD-based imaging system.

### Lifespan determination

*me*^*4J*^*/me*^*4J*^ mice were monitored from birth for survival and disease endpoint. Humane endpoints were defined as weight loss exceeding 20% of peak body weight, inability to ambulate, or severe footpad lesion causing distress. Mice reaching any humane endpoint were euthanized and recorded as deceased for Kaplan-Meier survival analysis. Survival curves were compared by log-rank (Mantel-Cox) test.

### Histology

Footpad and organ tissues were dissected and immediately fixed in 10% neutral buffered formalin for 24–48 hours. Fixed tissues were paraffin-embedded, sectioned at 5 μm, and stained with hematoxylin and eosin (H&E). For neutrophil enumeration, sections were stained with Ly6G or myeloperoxidase (MPO). Stained sections were imaged on a bright-field microscope at 10× and 40× magnification. Neutrophil infiltration was quantified by blinded counting of positive cells per high-power field (HPF) across at least five HPFs per section.

### Flow cytometry

Bone marrow, peripheral blood, footpad digests, or *in vitro* culture samples were processed into single-cell suspensions and stained with the antibody panels indicated in the Key Resources Table. For bone marrow preparation, femurs were collected, briefly sterilized in 70% ethanol, and marrow was flushed through a 70 μm strainer into FACS buffer using a 25G needle. After centrifugation, erythrocytes were lysed in ACK buffer for 10 minutes at room temperature, and cells were washed three times in PBS. Surface staining was performed in PBS containing 2% FBS for 30 minutes at 4°C. For intracellular phosphoprotein detection, cells were fixed in 4% paraformaldehyde for 10 minutes at 37°C, permeabilized in ice-cold 90% methanol for 30 minutes, and stained with anti-pERK1/2 overnight at 4°C. Data were acquired on an Attune NXT Acoustic Focusing Cytometer (Thermo Fisher) and analyzed with FlowJo (BD Biosciences).

For isolation of footpad-derived immune cells, tissues were kept on ice in FACS buffer until all samples had been collected. Samples were then incubated in enzymatic dissociation buffer containing collagenase D (1 mg) and DNase I (10 μL) in 10 mL FACS buffer at 37°C for up to 3 hours, followed by mechanical dissociation with a gentleMACS Dissociator (Miltenyi) using the Multi_H program for three cycles. Homogenates were filtered through a 70 μm strainer, centrifuged at 300g for 3 minutes at 4°C, resuspended in TrypLE, and incubated for 20 minutes at room temperature. After a second filtration and wash, cells were resuspended in FACS buffer and maintained on ice until staining. For surface staining, aliquots were transferred to microcentrifuge tubes, incubated with antibody panels for 1 hour at 4°C in the dark, washed with FACS buffer, and resuspended for acquisition. Where indicated, 200 μL aliquots were distributed in triplicate from each sample after the final resuspension step.

### pERK1/2 flow cytometry

Cells were stimulated in 1 mL medium with IL-1α alone, IL-15 alone, or IL-1α plus IL-15 at 10 ng/mL; additional experiments were performed at 30 and 50 ng/mL using the same design. Samples were incubated at 37°C for 0, 30, 60, 90, 120, or 150 seconds, immediately chilled on ice, and fixed by addition of an equal volume of 8% paraformaldehyde to achieve a final concentration of 4% PFA. After fixation for 15 minutes at room temperature in the dark, cells were washed in PBS, permeabilized in 90% methanol for 10 minutes at 4°C, and stained with the indicated flow cytometry antibodies for 1 hour at room temperature in the dark. Samples were washed in FACS buffer and acquired on an Attune NXT Acoustic Focusing Cytometer (Thermo Fisher).

For analysis of resting and activated neutrophil states, cells were co-stimulated with IL-1α plus IL-15 at either 10 ng/mL each or 50 ng/mL each. Cells were plated at 1 mL per well in 6-well plates and incubated at 37°C for 4 or 6 hours. Samples were then fixed to a final concentration of 4% paraformaldehyde for 15 minutes at room temperature in the dark, washed three times in PBS, surface-stained with the indicated antibodies for 1 hour at 4°C, washed in FACS buffer, and acquired on an Attune NXT Acoustic Focusing Cytometer (Thermo Fisher).

### Cytokine ELISA

For *in vitro* cytokine assays, isolated bone marrow neutrophils (5 × 10^5 cells per well) were plated in 96-well plates and stimulated with recombinant cytokines (IL-1α, IL-1β, IL-15, IL-2, IL-4, IL-7, IL-9, or IL-21; concentrations indicated in the figure legends), alone or in combination, in the presence or absence of inhibitors as indicated. Supernatants were collected at the specified time points, and cytokine concentrations were measured by ELISA according to the manufacturer’s instructions. Experiments were performed with three biological replicates, each assayed in duplicate technical wells.

### RNA extraction and RNA-seq (Figure 5 and SF3 and SF4)

Neutrophils were isolated by FACS based on co-expression of CD11b and Ly6G. CD11b+Ly6G+ cells were sorted using the BD FACSAria II at the Jackson Laboratory Flow Cytometry Core Facility (Bar Harbor, ME) and collected in FACS buffer for RNA-seq. For RNA isolation, library preparation, and sequencing, cells were homogenized in TRIzol Reagent (Thermo Fisher Scientific) by needle shearing, and total RNA was subsequently isolated using the miRNeasy Mini Kit (Qiagen) following the manufacturer’s protocol, including the optional on-column DNase digestion step to remove residual genomic DNA. RNA concentration and purity were evaluated using a NanoDrop 2000 spectrophotometer (Thermo Scientific), and RNA integrity was assessed using the Total RNA Pico Bioanalyzer Assay on an Agilent 2100 Bioanalyzer (Agilent Technologies). Stranded mRNA sequencing libraries were constructed from qualified RNA samples using the KAPA mRNA HyperPrep Kit (Roche Sequencing and Life Science) according to the manufacturer’s protocol. Briefly, polyadenylated mRNA was enriched from total RNA using oligo-dT magnetic beads, followed by thermal fragmentation and first- and second-strand cDNA synthesis. Illumina-compatible adapters containing unique dual-index barcode sequences were ligated to each library, and the libraries were subsequently amplified by PCR. Library size distribution and quality were assessed using the D5000 ScreenTape System on an Agilent TapeStation (Agilent Technologies), and library concentration was quantified using the Qubit dsDNA High Sensitivity (HS) Assay on a Qubit Fluorometer (Thermo Fisher Scientific). Sequencing was performed on an Illumina NextSeq 500 instrument using the High Output Reagent Kit v2.5, generating 75 bp single-end reads. Reads were quality-filtered with FastP v0.23.2, aligned to the GRCm39 reference genome using STAR v2.7.11, and gene-level counts were generated by RSEM v1.3.3. Expected read counts per gene produced by RSEM were rounded to integer values, filtered to include only genes that have at least two samples within a sample group having a cpm > 1.0, and were passed to edgeR (v4.5.0) for differential expression analysis. The negative binomial conditional common likelihood was maximized to estimate a common dispersion value across all genes. Exact tests were used to elucidate statistical differences between the two sample groups of negative binomially distributed counts producing p-values per test. The Benjamini and Hochberg’s algorithm (p-value adjustment) was used to control the false discovery rate (FDR). Features with an FDR-adjusted p-value < 0.05 were declared significantly differentially expressed. Data were analyzed through the use of QIAGEN IPA (QIAGEN Inc., https://www.qiagenbioinformatics.com/products/ingenuity-pathway-analysis)(Krämer et al., 2013).

### RNA extraction and RNA-seq (Figure 2 and SF1)

Neutrophils were isolated using the Miltenyi Mouse Neutrophil Isolation Kit according to the manufacturer’s protocol. Briefly, femurs were flushed with FACS buffer, filtered through a 70 μm cell strainer, and pelleted by centrifugation. Erythrocytes were lysed in ACK buffer for 10 minutes at room temperature, and cells were washed three times in PBS. Cells were resuspended in neutrophil isolation buffer (PBS containing 1% BSA and 2 mM EDTA) at the manufacturer’s recommended volume relative to total input cell number, incubated sequentially with Neutrophil Biotin-Antibody Cocktail and Anti-Biotin MicroBeads, and enriched on LS columns in a MidiMACS Separator. The unlabeled flow-through containing enriched neutrophils was collected for downstream RNA-seq processing.

RNA-seq was performed by Plasmidsaurus. Neutrophils were harvested in Zymo DNA/RNA Shield and submitted at approximately 100,000 cells per sample in 60 μL. Total RNA was extracted by automated bead-based methods, and polyadenylated mRNA was captured using oligo-dT primers incorporating sample barcodes and unique molecular identifiers. Libraries were generated using the vendor’s 3’ end counting workflow, followed by single-end Illumina sequencing (∼90-bp reads) targeting approximately 10 million deduplicated reads per sample. Reads were quality filtered with FastP v0.24.0, aligned with STAR v2.7.11, deduplicated by UMI with UMICollapse v1.1.0, and quantified at the gene level with featureCounts (subread v2.1.1) using strand-specific counting across exons and 3’ UTRs. Differential expression analysis was performed with edgeR v4.0.16.

### Analysis of public human transcriptomic datasets

No primary human transcriptomic experiments were performed in this study. Analyses of human neutrophil and inflammatory tissue transcriptomes presented in Figure 1 were conducted exclusively by re-analysis of publicly available datasets deposited in the NCBI Gene Expression Omnibus (GEO). Two datasets were retrieved and analyzed. GSE103170 is an RNA-seq dataset of peripheral blood neutrophils from 3 patients with active systemic juvenile idiopathic arthritis (sJIA) compared with 3 healthy controls. Raw HTSeq count files were downloaded from GEO. Differential expression analysis was performed on the raw counts using DESeq2 (via PyDESeq2), which employs a negative binomial generalized linear model. GSE148027 is an Affymetrix Human Genome U133 Plus 2.0 Array (GPL570) dataset comprising 18 lesional and 15 non-lesional/healthy skin biopsies from patients with hidradenitis suppurativa (HS). Processed series matrix files containing normalized, log2-transformed expression values were used directly. Where multiple probes mapped to a single gene, the probe with the highest mean expression was retained. Differential expression analysis for the microarray data was performed using Welch’s independent t-test. For both datasets, p-values were adjusted for multiple comparisons using the Benjamini-Hochberg (BH) method to control the false discovery rate (FDR). Pathway gene sets for the IL-1, γc, and MEK/ERK signaling pathways and their downstream transcriptional targets were manually curated based on canonical pathway definitions from KEGG and the primary literature. A complete list of gene set members is provided in Supplementary Table S1. Gene set enrichment analysis (GSEA) was performed using the GSEApy Python library (Fang et al., 2023; v1.0.5), utilizing the prerank module. Genes were ranked by their test statistic (Wald statistic for RNA-seq; t-statistic for microarray) to account for both fold-change direction and statistical significance. GSEA was run with 1,000 permutations, a minimum gene set size of 3, and a reproducibility seed of 42. Pathways with an FDR q-value < 0.25 were considered significantly enriched. Pathway module scores were calculated as the mean log_2_ fold-change of the genes within each curated gene set, with error bars representing the standard error of the mean (SEM). Data visualization was performed using matplotlib (v3.7.1) and seaborn (v0.12.2) in Python 3.11.

### Analysis of public human proteomic and phospho-proteomic datasets

No primary human proteomic experiments were performed in this study. All proteomic analyses of human neutrophils presented in Figure 3 were conducted by re-analysis of publicly available datasets retrieved from the PRIDE/ProteomeXchange repository (https://www.ebi.ac.uk/pride).

#### Dataset selection and data acquisition

Datasets were selected based on the inclusion of primary human neutrophils and the availability of publicly accessible processed quantitative protein tables. Three datasets were used for re-analysis of baseline neutrophil proteomes. The primary healthy-donor reference was PXD0107013, a deep DIA dataset comprising 83 healthy donors and 98 patients with monogenic neutrophil disorders, from which the author-processed Spectronaut quantification report was analyzed directly. Additional healthy-donor references included PXD0445694, a DIA dataset of tissue-specific neutrophils from which the four resting blood neutrophil samples were extracted using the deposited Spectronaut report, and PXD0043525, a MaxQuant-based immune cell proteome atlas from which neutrophil-specific unique peptide counts were extracted (n = 4 resting neutrophil donors).

#### Target protein selection and detection scoring

Proteins of interest were selected a priori based on membership in key neutrophil signaling pathways, including the IL-1/TLR pathway (IL1R1, MYD88, IRAK1, IRAK4, TRAF6), the γc cytokine pathway (IL2RB, IL2RG, IL4RA, IL15, IL15RA, JAK3), the NFκB/STAT signaling axes (STAT5A, STAT5B, STAT3, NFKB1, RELA), and the MAPK/ERK pathway (MAP2K1, MAPK1, MAPK3). Gene nomenclature follows HGNC conventions. To evaluate the presence of these low-abundance signaling nodes across datasets, detection confidence was scored on a three-level ordinal scale using the deposited processed outputs at a 1% protein-level false discovery rate. A score of 0 indicated no detected signal in the relevant neutrophil samples. A score of 1 indicated detection with exactly one unique or stripped peptide in at least one sample within a dataset. A score of 2 indicated detection with two or more unique or stripped peptides in at least one sample. Re-analysis heatmaps display proteins achieving a score of at least 1 in one or more analyzed datasets.

#### Phospho-proteomic analysis

Phospho-proteomic data for primary human neutrophils were extracted from dataset PXD029046. This study profiled global phosphorylation dynamics in neutrophils stimulated with LPS (2 μg/mL) and homocysteine (250 μM) for 30 minutes using TiO2 enrichment followed by LC-MS/MS. Quantitative fold-change values and p-values relative to unstimulated controls were extracted from the study’s supplementary materials and published kinase enrichment analyses. Phosphorylation responses were evaluated across four signaling axes: IL-1R/TLR, NFκB, γc cytokine, and MEK/ERK. When multiple phospho-sites were reported for a given protein, the site with the largest stimulation-dependent increase was used. Where available, fold-change values supported by orthogonal validation in the original study were prioritized. Proteins not detected under a given stimulation condition were annotated as not detected (ND).

## Resource availability

### Lead Contact

Further information and requests for resources and reagents should be directed to and will be fulfilled by the lead contact, Vishnu Hosur.

## Materials availability

All mouse strains generated in this study will be made available upon request to the lead contact. All other mouse strains used are commercially available from The Jackson Laboratory (see Key Resources Table S2).

## Supplementary Materials

Fig. S1 to S6 for multiple supplementary figures.

Supplemental figure legends

Table S1

Table S2

## Acknowledgments

We gratefully acknowledge the contributions of The Jackson Laboratory core facilities for their expert assistance at Genome Technologies, Genetic Engineering Technologies, Flow Cytometry, and Histopathology and Microscopy Sciences, and Computational Sciences.

## Author contributions

KL, LA, JW, LDS, and VH designed the experiments. KL, LA, LMB performed the experiments. TMS and VH contributed to bioinformatic analyses and analyzed the data. KL and VH wrote the manuscript. All authors critically reviewed and approved the final manuscript.

## Data and materials availability

All data are available in the main text or the supplementary materials. Transcriptomic datasets analyzed in this study are publicly available under accession numbers GSE103170 and GSE148027. Phospho-proteomic datasets were accessed from PRIDE under accession numbers PXD010701, PXD044569, PXD004352, and PXD029046.

## Figures

**Supplemental Table 1**. Full gene lists for IL-1, γc, and MEK/ERK pathway modules used in transcriptomic analyses of sJIA (GSE103170) and hidradenitis suppurativa (GSE148027) datasets. Gene names, log_2_ fold-change values, and Benjamini-Hochberg-adjusted p values are provided for each pathway module and disease cohort.

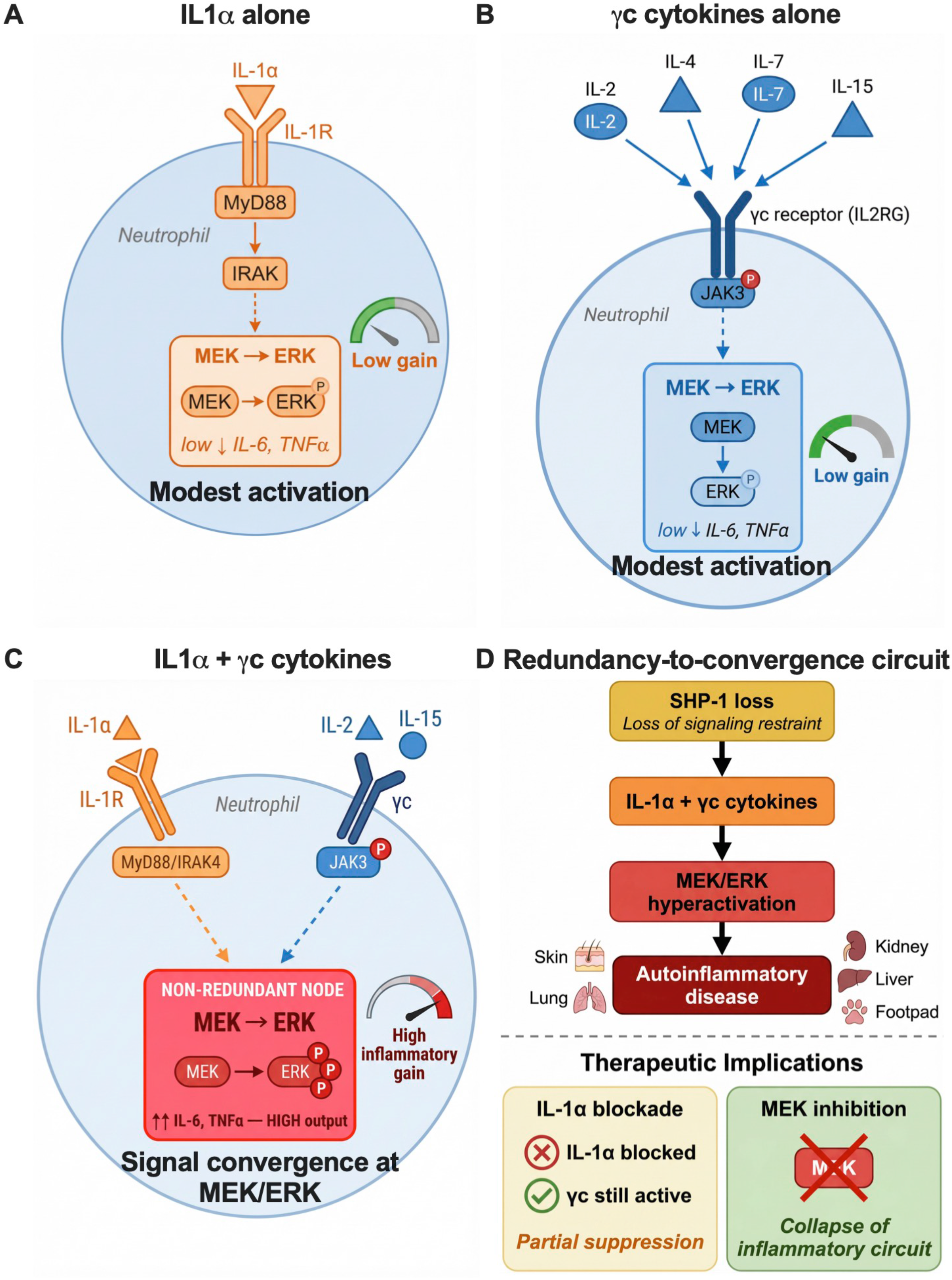

## References and Notes

1. A. Bertoni et al., A novel knock-in mouse model of cryopyrin-associated periodic syndromes with development of amyloidosis: Therapeutic efficacy of proton pump inhibitors. J Allergy Clin Immunol 145, 368–378.e313 (2020).

2. P. A. Nigrovic, P. Y. Lee, H. M. Hoffman, Monogenic autoinflammatory disorders: Conceptual overview, phenotype, and clinical approach. J Allergy Clin Immunol 146, 925–937 (2020).

3. E. Montaldo et al., Cellular and transcriptional dynamics of human neutrophils at steady state and upon stress. Nat Immunol 23, 1470–1483 (2022).

4. Y. Rochman, R. Spolski, W. J. Leonard, New insights into the regulation of T cells by γc family cytokines. Nature Reviews Immunology 9, 480–490 (2009).

5. C. L. Abram, C. A. Lowell, Shp1 function in myeloid cells. J Leukoc Biol 102, 657–675 (2017).

6. J. X. Lin, W. J. Leonard, The Common Cytokine Receptor γ Chain Family of Cytokines. Cold Spring Harb Perspect Biol 10, (2018).

7. A. B. Nesterovitch et al., Spontaneous insertion of a b2 element in the ptpn6 gene drives a systemic autoinflammatory disease in mice resembling neutrophilic dermatosis in humans. Am J Pathol 178, 1701–1714 (2011).

8. M. Speir, K. E. Lawlor, RIP-roaring inflammation: RIPK1 and RIPK3 driven NLRP3 inflammasome activation and autoinflammatory disease. Semin Cell Dev Biol 109, 114–124 (2021).

9. N. M. Ter Haar et al., Reversal of Sepsis-Like Features of Neutrophils by Interleukin-1 Blockade in Patients With Systemic-Onset Juvenile Idiopathic Arthritis. Arthritis Rheumatol 70, 943–956 (2018).

10. C. A. Penno et al., Lipidomics Profiling of Hidradenitis Suppurativa Skin Lesions Reveals Lipoxygenase Pathway Dysregulation and Accumulation of Proinflammatory Leukotriene B4. J Invest Dermatol 140, 2421–2432.e2410 (2020).

11. H. Ben Abdallah, A. Bregnhøj, L. Iversen, C. Johansen, Transcriptomic Analysis of Hidradenitis Suppurativa: A Unique Molecular Signature with Broad Immune Activation. Int J Mol Sci 24, (2023).

12. R. A. Brown et al., Neutrophils From Children With Systemic Juvenile Idiopathic Arthritis Exhibit Persistent Proinflammatory Activation Despite Long-Standing Clinically Inactive Disease. Front Immunol 9, 2995 (2018).

13. B. E. Rumberger, E. L. Boarder, S. L. Owens, M. D. Howell, Transcriptomic analysis of hidradenitis suppurativa skin suggests roles for multiple inflammatory pathways in disease pathogenesis. Inflamm Res 69, 967–973 (2020).

14. L. Mazgaeen et al., CD47 halts Ptpn6-deficient neutrophils from provoking lethal inflammation. Sci Adv 9, eade3942 (2023).

15. P. Y. Thimmappa et al., Quantitative phosphoproteomics reveals diverse stimuli activate distinct signaling pathways during neutrophil activation. Cell Tissue Res 389, 241–257 (2022).

16. N. D. Huntington et al., Interleukin 15-mediated survival of natural killer cells is determined by interactions among Bim, Noxa and Mcl-1. Nat Immunol 8, 856–863 (2007).

17. S. Tartey, P. Gurung, P. Samir, A. Burton, T. D. Kanneganti, Cutting Edge: Dysregulated CARD9 Signaling in Neutrophils Drives Inflammation in a Mouse Model of Neutrophilic Dermatoses. J Immunol 201, 1639–1644 (2018).

18. J. R. Lukens, T. D. Kanneganti, SHP-1 and IL-1α conspire to provoke neutrophilic dermatoses. Rare Dis 2, e27742 (2014).

19. A. T. Waickman, J. Y. Park, J. H. Park, The common γ-chain cytokine receptor: tricks-and-treats for T cells. Cell Mol Life Sci 73, 253–269 (2016).

20. V. Budagian, E. Bulanova, R. Paus, S. Bulfone-Paus, IL-15/IL-15 receptor biology: a guided tour through an expanding universe. Cytokine Growth Factor Rev 17, 259–280 (2006).

21. M. Pelletier, C. Ratthé, D. Girard, Mechanisms involved in interleukin-15-induced suppression of human neutrophil apoptosis: role of the anti-apoptotic Mcl-1 protein and several kinases including Janus kinase-2, p38 mitogen-activated protein kinase and extracellular signal-regulated kinases-1/2. FEBS Lett 532, 164–170 (2002).

22. C. Ratthé, D. Girard, Interleukin-15 enhances human neutrophil phagocytosis by a Syk-dependent mechanism: importance of the IL-15Ralpha chain. J Leukoc Biol 76, 162–168 (2004).

23. E. Bousoik, H. Montazeri Aliabadi, “Do We Know Jack” About JAK? A Closer Look at JAK/STAT Signaling Pathway. Front Oncol 8, 287 (2018).

24. M. Speir et al., Ptpn6 inhibits caspase-8-and Ripk3/Mlkl-dependent inflammation. Nature Immunology 21, 54–64 (2020).

25. T. Cai, R. Lenoir Capello, X. Pi, H. Wu, J. J. Chou, Structural basis of γ chain family receptor sharing at the membrane level. Science 381, 569–576 (2023).

26. E. A. Clark, L. D. Shultz, S. B. Pollack, Mutations in mice that influence natural killer (NK) cell activity.Immunogenetics 12, 601–613 (1981).

27. T. S. Migone et al., Recruitment of SH2-containing protein tyrosine phosphatase SHP-1 to the interleukin 2 receptor; loss of SHP-1 expression in human T-lymphotropic virus type I-transformed T cells. Proc Natl Acad Sci U S A 95, 3845–3850 (1998).

28. J. Y. Chu, I. Dransfield, A. G. Rossi, S. Vermeren, Non-canonical PI3K-Cdc42-Pak-Mek-Erk Signaling Promotes Immune-Complex-Induced Apoptosis in Human Neutrophils. Cell Rep 17, 374–386 (2016).

