## Supplementary figures and images for "Redundant γc cytokines license IL-1-driven neutrophil inflammation through MEK/ERK convergence"

### SF1

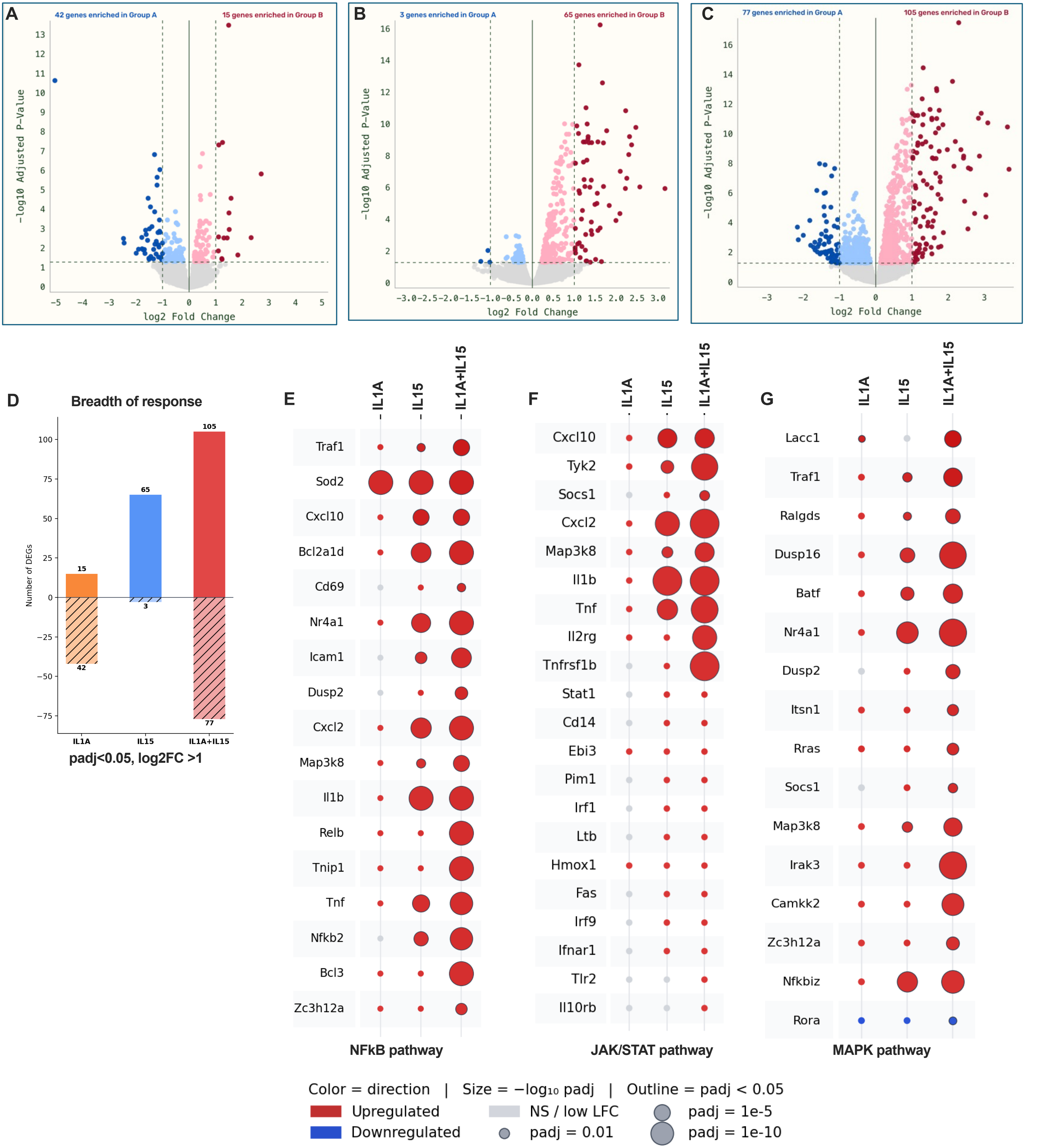

### SF2

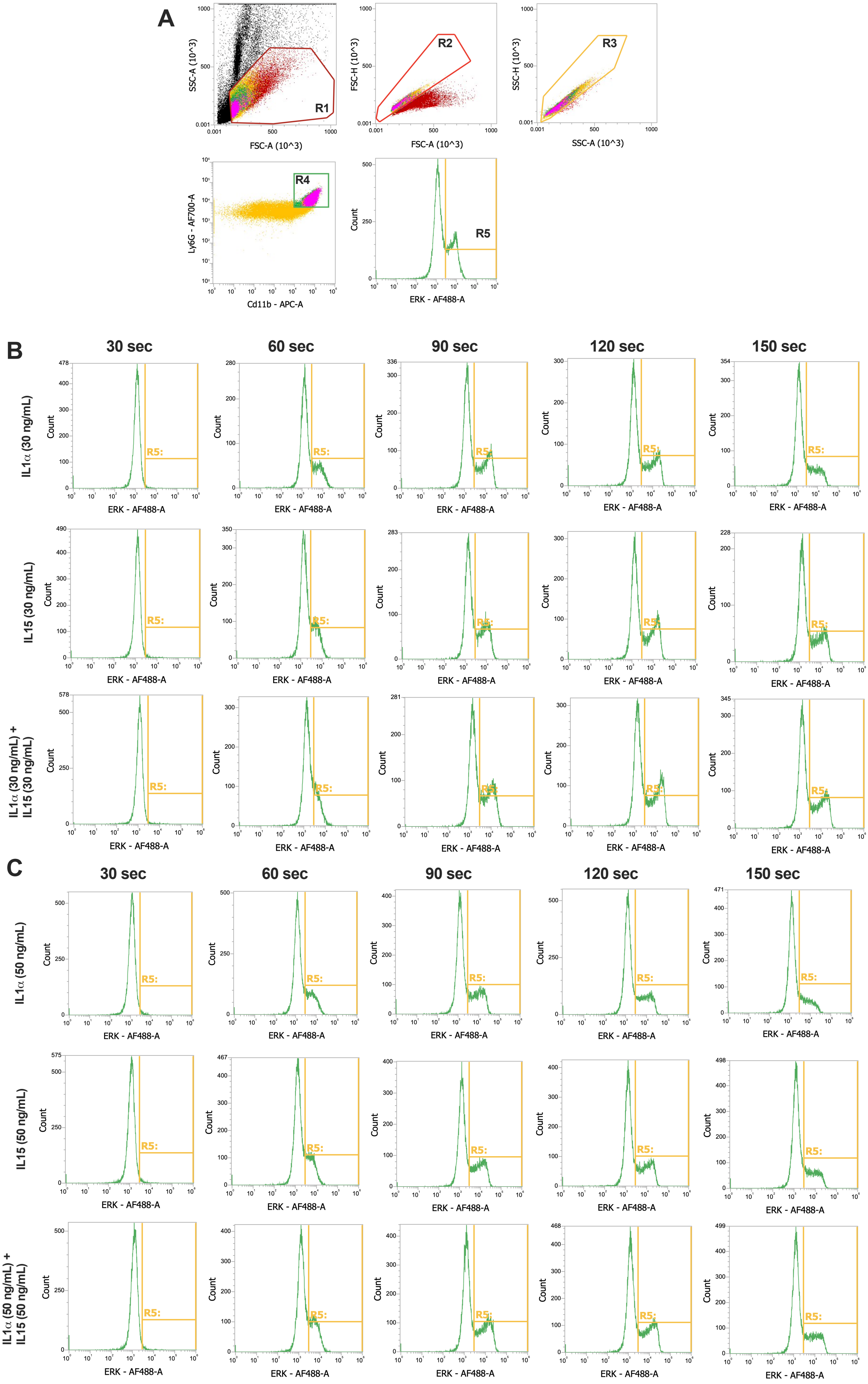

### SF3

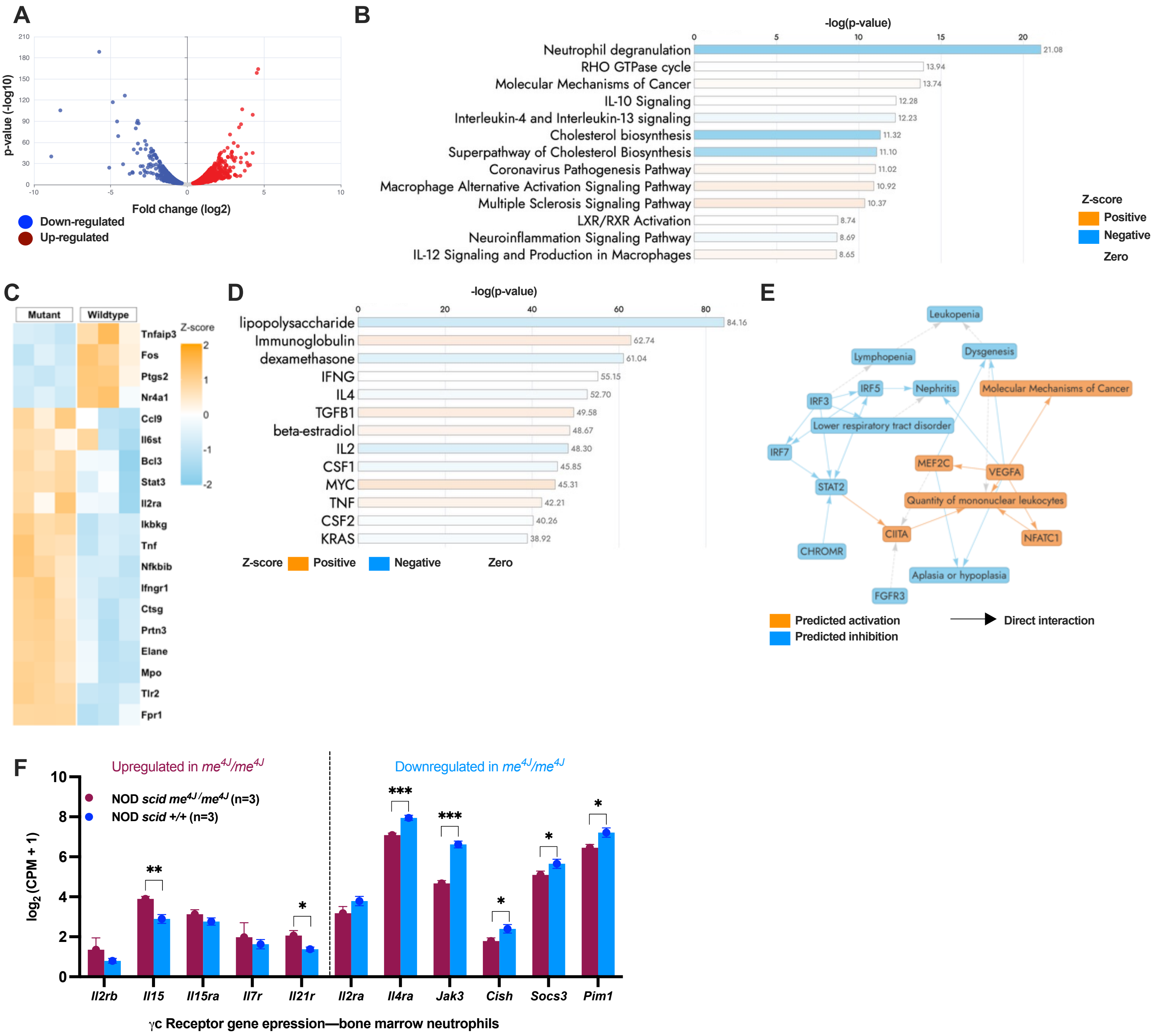

### SF4

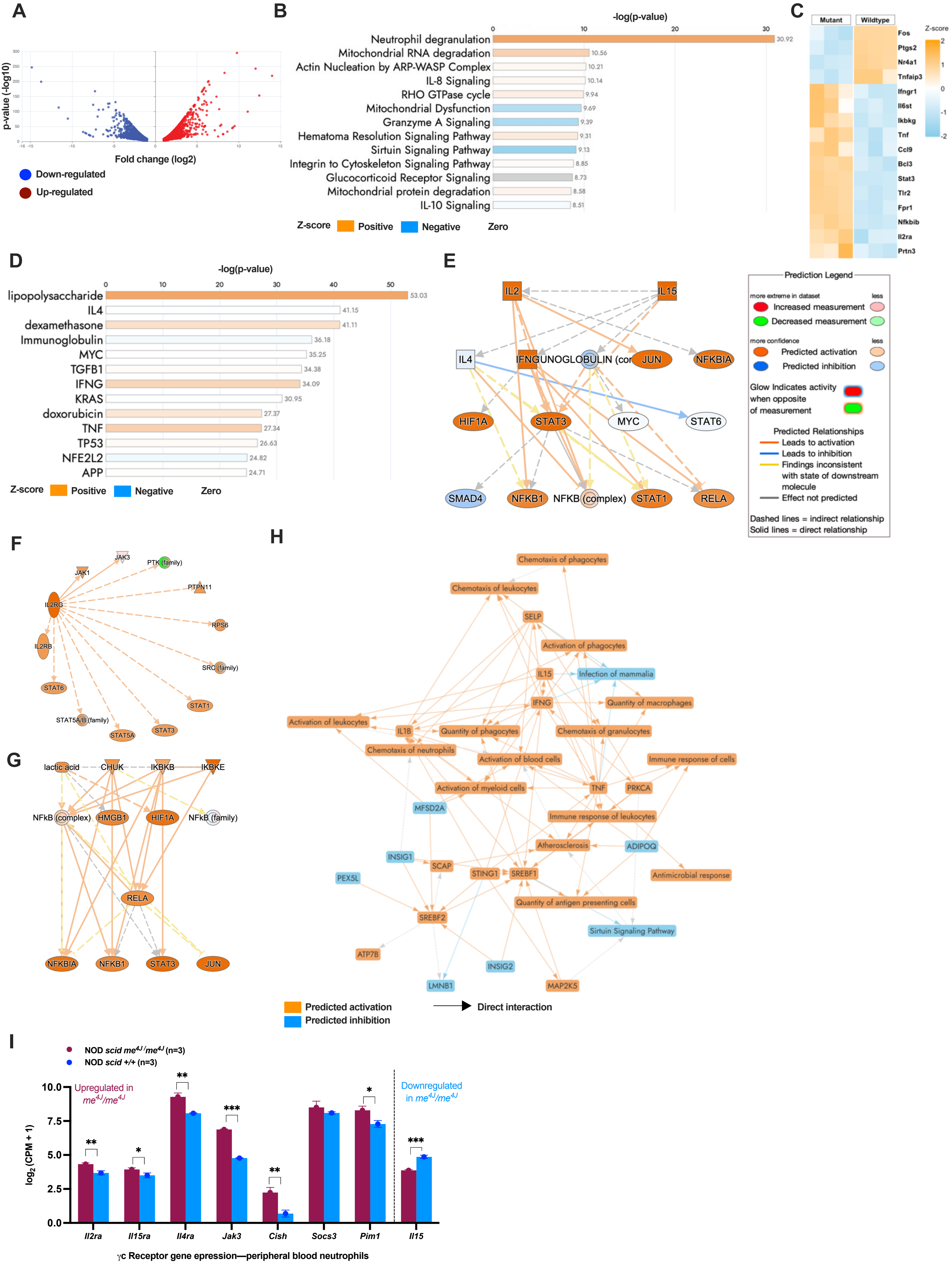

### SF5

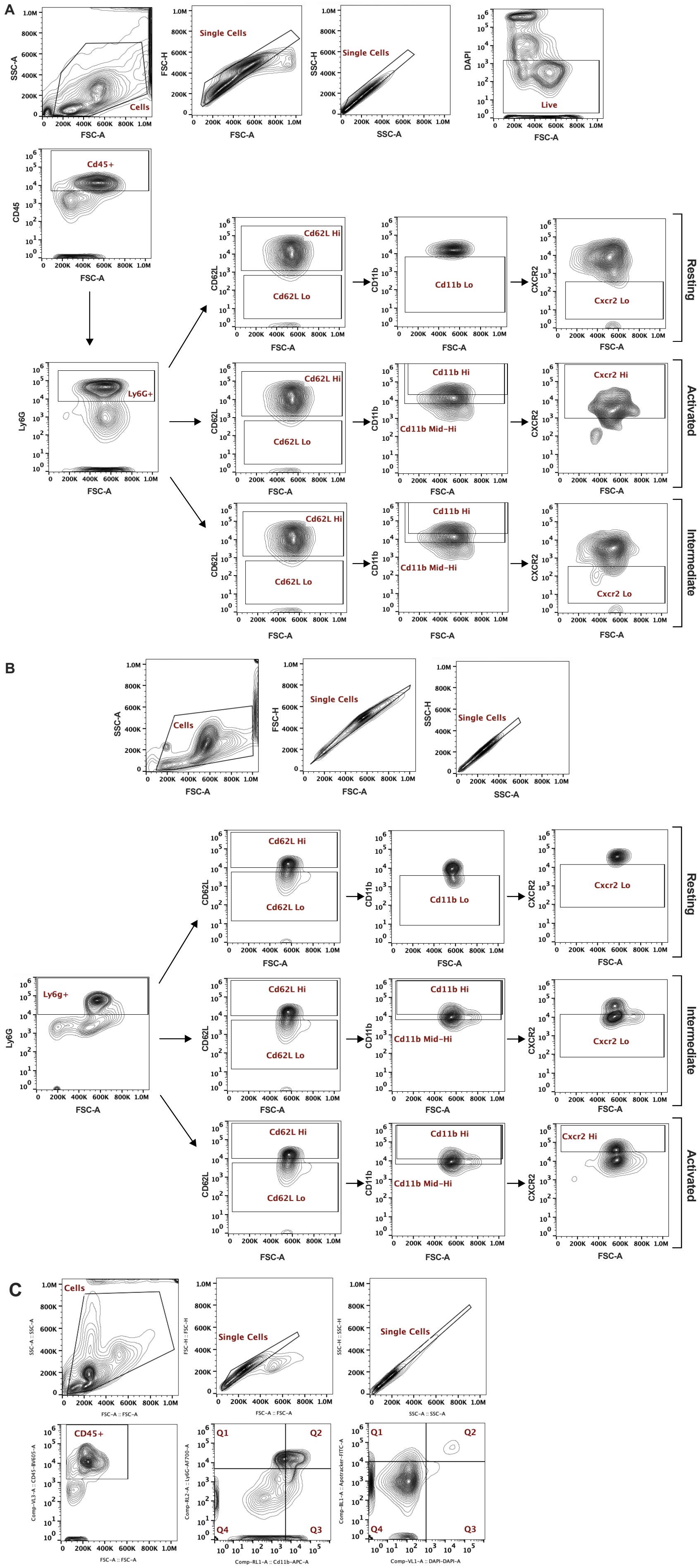
