## Supplementary material for "Redundant γc cytokines license IL-1-driven neutrophil inflammation through MEK/ERK convergence": SF6

**A** ***Il2rb***

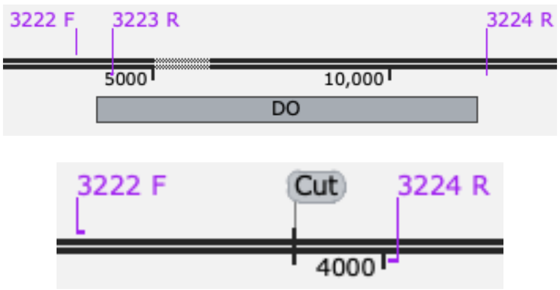


AGGGTTTGCATCCTCAGCTCCTCTCA
GCTGTGATGGCTACCATAGCTCTTGG
GCCATGGTAAGGAGACGCCTACCCA
GTCTTCCTTTCCCACCCAGTCTG

**B** ***Il4***

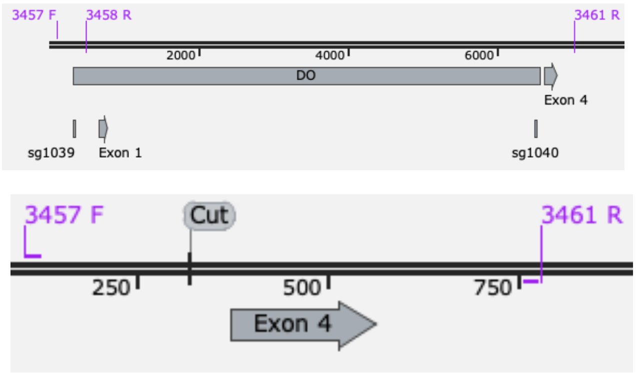


GACACCTGTGACCTCTTCCTTCTCTGCAGGAGGAGAGC
CAGTGGCAACCCAGGCCCGACAGCGAGACCCAAATCT
GTCTCACAATGAAACGTTTCTTTTA

**C** ***Il7r***

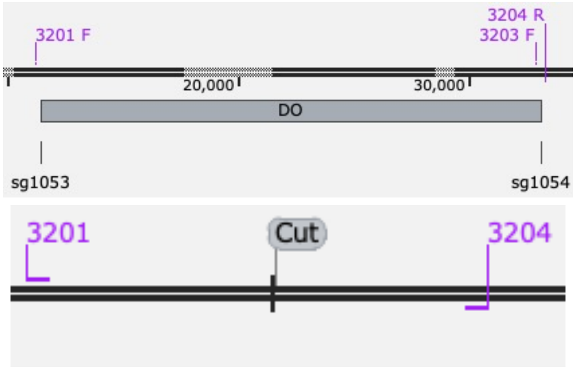


ACAGCTGTGTTTTGTTCTTCTCAGGAGACCTAGAAGATG
CAGACGCGGACCAGTTTCTCAGAGACAGCCCATCTCC
ACTTCCTCAGTACTGAATCAAGAA

**D** ***Il9r***

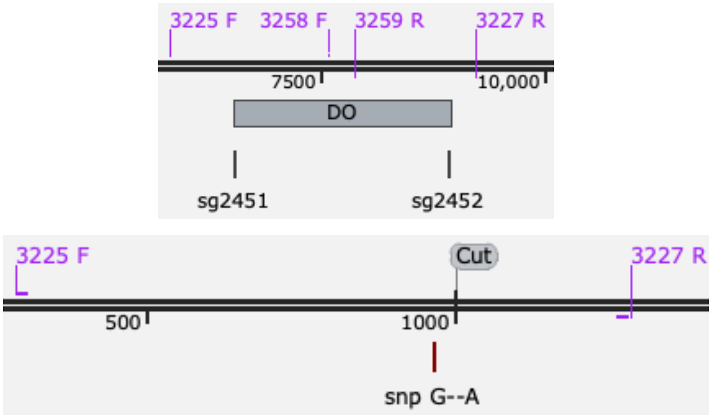


GACCTTGGAGAGAATGGCGGTGAAACAGGTCTCCTGGTT
CCTGATCTACATGCCCATCTTTCTTCTGCTGACTGGCTTTG
TCCACCTTCTGTTCAAGCTG

**E** ***Il15***

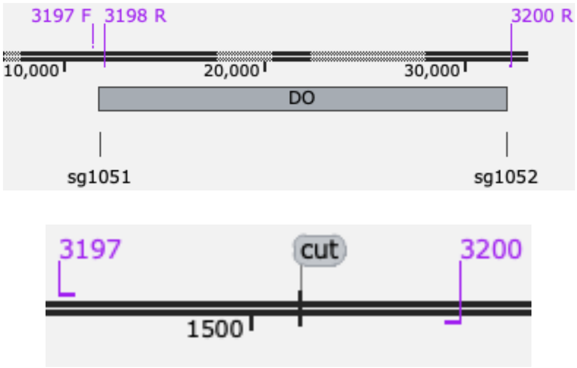


GTTTTACCATTTTGTCTACATTTATGTTTTCCAGAAACCA
TATATGAGGAGCTGGAGGAGAAAACCTTCACAGAGTT
TTTGCAAAGCTTTATACGCATTG

**F** ***Il21r***

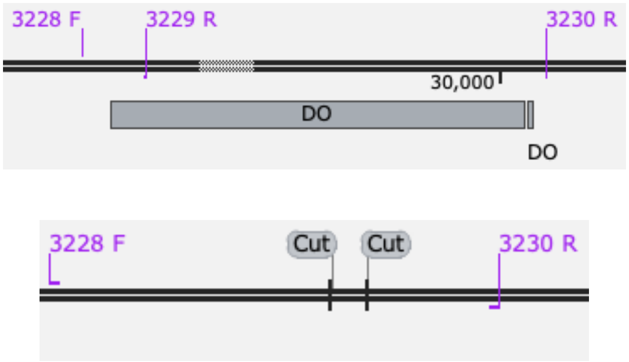


GTGCATGACCCCAGCTGGTTCACCACATATACCACATACT
TTTCTTGCAGGCTGGGACCCTCACATGCTGCTGCTCCTGG
CTGTCTTGATCATTGTCCTGGTTTTCATGGGTCTGAAGACC
TACTTTGGCCAGGGGACGTGGTGAGAGCACTGATGCTAAT
GAATAGT
