## Supplemental Figure Legends for "Redundant γc cytokines license IL-1-driven neutrophil inflammation through MEK/ERK convergence"

**Supplemental Figure and Table Legends**

**Supplemental Table 1.** Gene lists and statistical results for IL-1, γc cytokine, and MEK/ERK pathway analysis in human disease datasets. Gene lists used for pathway module scoring, GSEA, and heatmap analysis in Figure 1A–F. The table contains 80 genes organized into three pathway modules: IL-1 pathway (n=30), γc cytokine pathway (n=27), and MEK/ERK pathway (n=23). For each gene, statistical results are provided from two independent publicly available human disease datasets: sJIA neutrophils (GSE103170) and hidradenitis suppurativa (HS) skin (GSE148027). Columns include gene symbol, gene description, t-statistic, adjusted p-value (padj), and significance call for each dataset. Significance thresholds: *padj<0.05, **padj<0.01, ***padj<0.001; ns, not significant; n.d., not determined (gene not detected in dataset). Statistical comparisons represent disease versus control for each respective dataset.

**Supplemental Figure 1**. Combined IL-1α and IL-15 stimulation broadens the transcriptional inflammatory response through convergent activation of NFκB, JAK/STAT, and MAPK signaling pathways in bone marrow-derived neutrophils from NOD *scid* +/+ mice.

(A–C) Volcano plots showing differential gene expression in bone marrow-derived neutrophils from NOD *scid* +/+ mice stimulated for 3 hours with IL-1α (10 ng/mL; A), IL-15 (100 ng/mL; B), or IL-1α+IL-15 (10+100 ng/mL; C) versus unstimulated cells. The x-axis shows log_2_ fold change (Group B: stimulated / Group A: unstimulated) and the y-axis shows −log_10_ adjusted p-value. Dark red points indicate genes significantly enriched in the stimulated condition (Group B; log_2_FC>1, padj<0.05) and blue points indicate genes significantly enriched in the unstimulated condition (Group A; log_2_FC<−1, padj<0.05). The number of significantly enriched genes in each group is indicated above each plot. Differential expression analysis was performed using edgeR.

(D) Bar chart summarizing the number of differentially expressed genes (DEGs) per stimulation condition (padj<0.05, |log_2_FC| > 1). Solid bars indicate genes upregulated in the stimulated condition; hatched bars indicate genes downregulated in the stimulated condition relative to unstimulated cells.

(E–G) Dot plots showing log₂ fold change of selected genes from the NFκB (E), JAK/STAT (F), and MAPK (G) signaling pathways across IL-1α, IL-15, and IL-1α+IL-15 stimulation conditions versus unstimulated cells at 3 hours post-stimulation. Genes are sorted by IL-1α+IL-15 log_2_FC in descending order. Dot size reflects statistical significance (−log_10_ padj); dot color indicates direction of regulation (red, upregulated; blue, downregulated; grey, not significant or |log_2_FC| < 0.3); outlined dots indicate padj<0.05. NFκB pathway genes are activated primarily by IL-1α and progressively amplified by combined stimulation (E). JAK/STAT pathway genes are activated predominantly by IL-15 (F). MAPK pathway genes show preferential activation under combined IL-1α+IL-15 stimulation, consistent with the convergent MEK/ERK signaling described in Figure 2N (G).

**Supplemental Figure 2**. Synergistic ERK1/2 phosphorylation in bone marrow-derived neutrophils from NOD scid+/+ mice stimulated with IL-1α and IL-15.

(A) Gating strategy for identification and analysis of phospho-ERK1/2 (pERK1/2) in bone marrow-derived neutrophils. R1: viable cell scatter gate excluding debris; R2: singlet discrimination by FSC-H vs FSC-A; R3: singlet confirmation by SSC-H vs SSC-A; R4: neutrophil identification by Ly6G (AF700) and CD11b (APC) co-expression; R5 (yellow gate): pERK1/2-positive population defined against an unstimulated control.

(B) Representative intracellular flow cytometry histograms showing pERK1/2 (ERK-AF488) signal in bone marrow-derived neutrophils from NOD *scid* +/+ mice stimulated with IL-1α (30 ng/mL), IL-15 (30 ng/mL), or IL-1α+IL-15 (30+30 ng/mL) at 30, 60, 90, 120, and 150 seconds post-stimulation. The R5 gate (yellow) demarcates the pERK1/2-positive population defined relative to unstimulated cells.

(C) Representative intracellular flow cytometry histograms as in (B) from an independent experiment using higher doses of IL-1α (50 ng/mL), IL-15 (50 ng/mL), or IL-1α+IL-15 (50+50 ng/mL). Combined IL-1α+IL-15 stimulation produces a reproducible rightward shift in pERK1/2 signal and an increase in the R5-gated pERK1/2-positive population compared to either cytokine alone across both doses and all timepoints, indicating synergistic ERK1/2 activation within 30–150 seconds of stimulation.

**Supplemental Figure 3**. Transcriptional profiling of bone marrow-derived neutrophils from NOD *scid-me^4J^/me^4J^* and NOD *scid* +/+ mice.

(A) Volcano plot of differentially expressed genes (DEGs) in bone marrow-derived neutrophils isolated from NOD *scid*-*me^4J^/me^4J^* versus NOD *scid* +/+ mice. Each point represents one gene; upregulated genes are shown in red and downregulated genes in blue. The x-axis represents log_2_ fold change, and the y-axis represents statistical significance (−log_10_ p-value).

(B) IPA canonical pathway analysis of DEGs ranked by −log_10_ (p-value), with values shown to the right of each bar. Bar color indicates predicted activation state: orange, predicted activation (positive Z-score); blue, predicted inhibition (negative Z-score); white, no directional prediction. Neutrophil degranulation was the most significantly enriched pathway (−log_10_ (p) = 21.08), followed by RHO GTPase cycle, Molecular Mechanisms of Cancer, IL-10 Signaling, and Interleukin-4 and Interleukin-13 signaling.

(C) Heatmap of selected inflammation-associated DEGs in bone marrow-derived neutrophils. Each column represents one biological replicate (Mutant: NOD *scid*-*me^4J^/me^4J^*, n=3; Wildtype: NOD scid +/+, n=3). Color scale represents Z-score normalized expression (orange, high; blue, low). The upper cluster contains genes upregulated in NOD *scid*-*me^4J^/me^4J^* (Tnfaip3, Fos, Ptgs2, Nr4a1); the lower cluster contains genes downregulated in NOD *scid*-*me^4J^/me^4J^* (Ccl9, Il6st, Bcl3, Stat3, Il2ra, Ikbkg, Tnf, Nfkbib, Ifngr1, Ctsg, Prtn3, Elane, Mpo, Tlr2, Fpr1).

(D) IPA upstream regulator analysis of DEGs ranked by −log_10_ (p-value). Bar color indicates predicted activation state of each regulator. LPS was the most significant predicted upstream regulator (−log_10_ (p) = 84.16), followed by immunoglobulin, dexamethasone, IFNG, IL4, TGFB1, beta-estradiol, IL2, CSF1, MYC, TNF, CSF2, and KRAS.

(E) IPA-generated network summarizing predicted relationships between key upstream regulators and downstream biological outcomes. Orange nodes indicate predicted activation; blue nodes indicate predicted inhibition; solid arrows represent direct interactions. Key predicted downstream outcomes include dysgenesis, leukopenia, lymphopenia, nephritis, lower respiratory tract disorder, and altered quantity of mononuclear leukocytes.

(F) Expression of γc cytokine receptor subunit and downstream signaling genes in bone marrow-derived neutrophils from NOD *scid*-*me^4J^/me^4J^* (dark red, n=3) and NOD *scid* +/+ (blue, n=3) mice. Bar height represents mean expression (log_2_[CPM+1]); error bars indicate standard deviation; individual data points are overlaid. Genes are grouped by direction of differential expression: upregulated in NOD *scid*-*me^4J^/me^4J^* (*Il2rb, Il15, Il15ra, Il7r, Il21r*; left of dashed line) and downregulated in NOD *scid*-*me^4J^/me^4J^* (*Il2ra, Il4ra, Jak3, Cish, Socs3, Pim1*; right of dashed line). Statistical significance was determined by unpaired two-tailed Student's t-test; *p<0.05, **p<0.01, ***p<0.001.

**Supplemental Figure 4**. **Transcriptional profiling of peripheral blood neutrophils from NOD *scid*-*me^4J^/me^4J^* and NOD *scid* +/+ mice.**

(A) Volcano plot of differentially expressed genes (DEGs) in peripheral blood neutrophils from NOD *scid*-*me^4J^/me^4J^* versus NOD *scid* +/+ mice. Upregulated genes are shown in red and downregulated genes in blue. The x-axis represents log₂ fold change, and the y-axis represents statistical significance (−log_10_ p-value). Peripheral blood neutrophils displayed larger fold changes (up to ±15 log_2_FC) compared to bone marrow neutrophils (±10 log₂FC, Supplemental Figure 3A), indicating a more pronounced transcriptional phenotype in circulating neutrophils.

(B) IPA canonical pathway analysis of DEGs from peripheral blood neutrophils ranked by −log_10_ (p-value). Bar color indicates predicted activation state: orange, predicted activation; blue, predicted inhibition; grey, no directional prediction. Neutrophil degranulation was the most significantly enriched pathway (−log_10_ (p) = 30.92), followed by Mitochondrial RNA degradation, Actin Nucleation by ARP-WASP Complex, IL-8 Signaling, RHO GTPase cycle, Mitochondrial Dysfunction, Granzyme A Signaling, Hematoma Resolution Signaling Pathway, and Sirtuin Signaling Pathway.

(C) Heatmap of selected inflammation-associated DEGs in peripheral blood neutrophils. Each column represents one biological replicate (Mutant: NOD *scid*-*me^4J^/me^4J^*, n=3; Wildtype: NOD scid +/+, n=3). Color scale represents Z-score normalized expression (orange, high; blue, low). The upper cluster contains genes upregulated in NOD *scid*-*me^4J^/me^4J^* (Fos, Ptgs2, Nr4a1, Tnfaip3); the lower cluster contains genes downregulated in NOD *scid*-*me^4J^/me^4J^* (Ifngr1, Il6st, Ikbkg, Tnf, Ccl9, Bcl3, Stat3, Tlr2, Fpr1, Nfkbib, Il2ra, Prtn3).

(D) IPA upstream regulator analysis ranked by −log_10_(p-value). LPS was the most significant predicted upstream regulator (−log_10_(p) = 53.03), followed by IL4, dexamethasone, immunoglobulin, MYC, TGFB1, IFNG, KRAS, doxorubicin, TNF, TP53, NFE2L2, and APP.

(E) IPA-generated network showing predicted activation of IL-15 and IL-2 signaling in peripheral blood neutrophils. IL-15 and IL-2 are predicted upstream activators driving STAT1, STAT3, NFKB1, RELA, JUN, and NFKBIA. Orange = predicted activation; blue = predicted inhibition. Solid lines = direct relationships; dashed lines = indirect.

(F) IPA-generated network showing IL2RG (γc) as a central hub activating JAK1, JAK3, IL2RB, STAT1, STAT3, STAT5A, STAT5A/B, and STAT6.

(G) IPA-generated network showing IKBKE/NFκB signaling axis activation. IKBKE, IKBKB, and CHUK converge on the NFκB complex, HMGB1, and HIF1A, driving downstream RELA, NFKB1, STAT3, and JUN.

(H) IPA-generated graphical summary network integrating predicted upstream activators with downstream biological functions. Predicted activated functions (orange): chemotaxis of phagocytes and leukocytes, activation of phagocytes, myeloid cells, and blood cells, immune response of leukocytes, antimicrobial response. Predicted inhibited nodes (blue): infection of mammalia, atherosclerosis, sirtuin signaling pathway. Solid arrows = direct; dashed arrows = indirect.

(I) Expression of γc cytokine receptor subunit and downstream signaling genes in peripheral blood neutrophils from NOD *scid*-*me^4J^/me^4J^* (dark red, n=3) and NOD *scid* +/+ (blue, n=3) mice. Bar height = mean expression (log_2_[CPM+1]); error bars = SD; individual data points overlaid. Upregulated in NOD *scid*-*me^4J^/me^4J^* (*Il2ra, Il15ra, Il4ra, Jak3, Cish, Socs3, Pim1*; left of dashed line); downregulated (Il15; right of dashed line). Note that directionality of *Jak3, Cish, Il15* is opposite to that in bone marrow neutrophils (Supplemental Figure 3F), indicating compartment-specific transcriptional reprogramming. Statistical significance by unpaired two-tailed Student's t-test; *p<0.05, **p<0.01, ***p<0.001.

**Supplemental Figure 5**. **Gating strategy for neutrophil activation state and apoptosis assays.**

**(A)** Representative gating strategy for *in vivo* neutrophil activation state analysis. Cells were sequentially gated on: FSC-A/SSC-A (Cells) → FSC-H/FSC-A single cells → SSC-H/SSC-A single cells → DAPI⁻ (Live) → CD45⁺ → Ly6G⁺. Ly6G⁺CD45⁺ neutrophils were then classified into three activation states based on sequential CD62L, CD11b, and CXCR2 expression: Resting (CD62L^hi^ CD11b^lo^ CXCR2^lo^), Intermediate (CD62L^lo^ CD11b^mid-hi^ CXCR2^lo^), and Activated (CD62L^lo^ CD11b^hi^ CXCR2^hi^). This gating strategy was applied to *in vivo* samples from NOD *scid*-*me^4J^/me^4J^* and NSG-*me^4J^/me^4J^* mice as shown in Figures 6A–6D.

**(B)** Representative gating strategy for *in vitro* stimulated bone marrow neutrophils (corresponding to Figures 6E–6G). Cells were gated on: FSC-A/SSC-A (Cells) → FSC-H/FSC-A single cells → SSC-H/SSC-A single cells → Ly6G⁺. The same CD62L → CD11b → CXCR2 sequential gating scheme was applied to classify neutrophils into Resting, Intermediate, and Activated populations, as defined in (A).

**(C)** Representative gating strategy for the ApoTracker apoptosis assay (corresponding to Figures 6H, 6M, and 6N). Cells were gated on: FSC-A/SSC-A (Cells) → FSC-H/FSC-A single cells → SSC-H/SSC-A single cells → CD45⁺. ApoTracker-FITC⁺DAPI⁻ cells (Q2) were scored as apoptotic neutrophils.

**Supplemental Figure 6.** CRISPR/Cas9-mediated sequence drop-out strategy at six cytokine receptor loci.

Schematic diagrams and validation sequences for CRISPR/Cas9-based KO mice generated at the *Il2rb* (A), *Il4* (B), *Il7r* (C), *Il9r* (D), *Il15* (E), and *Il21r* (F) loci. For each locus, the upper panel shows the genomic region with the dropped-out sequence (DO), sgRNA cut site(s), and flanking PCR primer locations (purple, numbered). The lower panel shows a zoomed view of the cut site with genotyping primers. Sequences below each diagram display the portion of the PCR amplicon directly flanking each cut site. Mutant sequences are shown in red. For *Il9r* (D), a G-to-A SNP at the junction is indicated. For *Il21r* (F), two cut sites flank two independent drop-out regions; black sequence indicates the common sequence upstream of the first dropout,

red indicates the WT sequence retained between the two dropouts, and blue indicates the common sequence downstream of the second dropout. Mice were generated by zygotic electroporation of Cas9 and sgRNA(s)^1^.

**References**

1.	Hosur, V., Low, B.E., and Wiles, M.V. (2024). Chapter 18 - Genetic modification of mice using CRISPR-Cas9: Best practices and practical concepts explained. In Rigor and Reproducibility in Genetics and Genomics, D.F. Dluzen, and M.H.M. Schmidt, eds. (Academic Press), pp. 425-452. <https://doi.org/10.1016/B978-0-12-817218-6.00018-8>.
