## Supplementary material for "Redundant γc cytokines license IL-1-driven neutrophil inflammation through MEK/ERK convergence": Table S2

**Key Resources Table 2**

| **REAGENT or RESOURCE** | **SOURCE** | **IDENTIFIER** |
| --- | --- | --- |
| ***Antibodies*** | | |
| Rabbit anti-mouse PTPN6 mAb | Abcam | Cat # ab32559 |
| Rabbit anti-mouse GAPDH mAb | Cell Signaling Technology | Cat # 2118; RRID: AB_561053 |
| HRP-conjugated anti-rabbit IgG secondary antibody | Cell Signaling Technology | Cat # 7074; RRID: AB_2099233 |
| Anti-Ly6G (clone 1A8) | BD Biosciences | Cat # 551461; RRID: AB_394204 |
| Alexa Fluorophore 488 anti-ERK1/2 Phospho (Thr202/Tyr204) (clone 4B11B69) | BioLegend | Cat # 675508 |
| APC anti-mouse/human CD11b (clone M1/70) | BioLegend | Cat # 101212 |
| Alexa Fluorophore 700 anti-mouse Ly6G (clone 1A8) | BioLegend | Cat # 127622 |
| PE/Cyanine7 anti-mouse CD182 (CXCR2) (clone SA044G4) | BioLegend | Cat # 149316 |
| Brilliant Violet 605 anti-mouse CD184 (CXCR4) (clone L276F12) | BioLegend | Cat # 146519 |
| Brilliant Violet 605 anti-mouse CD45 (clone 30-F11) | BioLegend | Cat # 103139 |
| Apotracker Green | BioLegend | Cat # 427402 |
| BUV737 Rat Anti-Mouse CD45 (clone 30-F11) | BD Biosciences | Cat # 568344 |
| CD11b FITC | BD Biosciences | Cat # 564454 |
| CD62L BV711 | BD Biosciences | Cat # 568286 |
| ***Bacterial and Virus Strains*** | N/A |  |
| ***Biological Samples*** |  |  |
| sJIA peripheral blood neutrophil RNA-seq dataset | [GEO: GSE103170](Ter Haar et al., 2018) | GEO: GSE103170 |
| Hidradenitis suppurativa lesional skin transcriptomic dataset | [GEO: GSE148027](Penno et al., 2020) | GEO: GSE148027 |
| Human neutrophil proteome — healthy donors and monogenic neutrophil disorders | [PRIDE: PXD010701](Grabowski et al., 2019) | ProteomeXchange: PXD010701 |
| Human neutrophil proteome — absolute copy number quantification | [PRIDE: PXD044569](Sollberger et al., 2024) | ProteomeXchange: PXD044569 |
| Immune-cell proteome atlas (neutrophil fraction) | [PRIDE: PXD004352](Rieckmann et al., 2017) | ProteomeXchange: PXD004352 |
| Human neutrophil phospho-proteome — LPS, high glucose, homocysteine stimulation | [PRIDE: PXD029046](Thimmappa et al., 2022) | ProteomeXchange: PXD029046 |
| ***Chemicals, Peptides, and Recombinant Proteins*** |  |  |
| Recombinant murine IL-1α | BioLegend | Cat # 575006 |
| Recombinant murine IL-1β | BioLegend | Cat # 575102 |
| Recombinant murine IL-15 | BioLegend | Cat # 575104 |
| Recombinant murine IL-2 | BioLegend | Cat # 575404 |
| Recombinant murine IL-4 | BioLegend | Cat # 574302 |
| Recombinant murine IL-7 | BioLegend | Cat # 577802 |
| Recombinant murine IL-9 | BioLegend | Cat # 556004 |
| Recombinant murine IL-21 | BioLegend | Cat # 574504 |
| Trametinib (MEK inhibitor) | MedChemExpress | Cat # HY-10999 |
| Zimlovisertib (IRAK4 inhibitor) | MedChemExpress | Cat # HY-19836 / synonym:  PF-06650833 |
| Capivasertib (AKT inhibitor) | MedChemExpress | Cat # HY-15431 |
| Ritlecitinib (JAK3 inhibitor) | MedChemExpress | Cat # HY-100754 |
| AC-4-130 (STAT5 inhibitor) | MedChemExpress | Cat # HY-124500 |
| Igepal CA-630 (NP-40 substitute) | Sigma-Aldrich | Cat # I8896 |
| cOmplete protease inhibitor cocktail | Roche Diagnostics | Cat # 11836153001 |
| Collagenase D | Sigma-Aldrich | Cat # 11088866001 |
| DNase I | New England Biolabs | Cat # M0303L |
| TrypLE™ Express Enzyme | ThermoFisher Scientific | Cat # 12604013 |
| RBC Lysis Buffer (10X) | BioLegend | Cat # 420301 |
| DNA/RNA Shield | ZymoResearch | Cat # R1100-50 |
| Staurosporine | Tocris | Cat # 1285 |
| Phusion high-fidelity DNA polymerase | NEB | M0530L |
| QIAquick Gel Extraction Kit | Qiagen | Cat # 28704 |
| Mag beads PCR Product Cleanup Reagent | New England Biolabs | Cat # T4130S |
| LS column | Miltenyi Biotec | Cat # 130-042-401 |
| MidiMACS Separator | Miltenyi Biotec | Cat # 130-042-301 |
| 4–12% Tris-glycine PAGEr Gold Precast Gel | Lonza | Cat # LZ58100 |
| West Pico PLUS Chemiluminescent Substrate | Pierce/Thermo Fisher | Cat # 34577 |
| 5% non-fat dry milk in TBS-T | Bio-Rad | Cat # 1706404 |
| ***Critical Commercial Assays/kits*** |  |  |
| Mouse IL-6 ELISA | R&D systems | Cat # DY406 |
| Mouse TNFα ELISA | R&D systems | Cat # DY410 |
| Mouse G-CSF ELISA | R&D Systems | Cat # DY414 |
| Mouse GM-CSF ELISA | R&D Systems | Cat # DY415 |
| Mouse CXCL1/KC ELISA | R&D Systems | Ca t# DY453 |
| Mouse IL-1β/IL-1F2 ELISA | R&D Systems | Cat # DY401 |
| Neutrophil Isolation kit, Mouse | Miltenyi Biotec | Cat # 130-097-658 |
| Advantage GC 2 PCR kit | Takara Bio | Cat # 639119 |
| ExoSAP-IT™ PCR Product Cleanup Reagent | Thermo Fisher Scientific | Cat # 78205.10.ML |
| BigDye™ Terminator v3.1 Cycle Sequencing Kit | Thermo Fisher Scientific | Cat # 4337455 |
| ***Deposited Data*** |  |  |
| RNA-seq data | 19NGS_007_TED_GEO_04072026  19NGS_008_TED_GEO_04072026  Plasmidsaurus_GEO_040702026 | GSE327729  GSE327730  GSE327728 |
| ***Experimental Models: Organisms/Strains*** |  |  |
| NOD *scid-me^4J^/me^4J^* | JAX | stock: 41651 |
| NSG-*me^4J^/me^4J^* | JAX | stock: 41652 |
| NOD *scid* IL-15 Tg/Tg | JAX | stock: 39129 |
| NOD *scid Il2rb* KO | JAX | stock: 39085 |
| NOD *scid Il15* KO | JAX | stock: 40250 |
| NOD *scid Il4* KO | JAX | stock: 40251 |
| NOD *scid Il7r* KO | JAX | stock: 39465 |
| NOD *scid Il9r* KO | JAX | stock: 39087 |
| NOD *scid Il21r* KO | JAX | stock: 39086 |
| ***Oligonucleotides*** |  |  |
| 2955 F | *Ptpn6* | GAGGTCACCTTGCCTTCTTTAT |
| 2955 R | *Ptpn6* | GGCAGGGATAAATGTGTCTAGG |
| 3197 F | *Il15* | GGGCAAGAGGAAGAGGATAATG |
| 3198 R | *Il15* | GGTGCCGAAATCCTCAGTTA |
| 3200 R | *Il15* | TGTGATCCAAGTGGCTCATTAT |
| 3201 F | *Il7r* | ACAGAGAGAGCACATTGAAACT |
| 3203 F | *Il7r* | GCCCACCAGAAACAGTTAGA |
| 3204 R | *Il7r* | GGTCACGTTGACTTTCTCTTCT |
| 3222 F | *Il2rb* | GTGGATGCTTCTCGCCTATT |
| 3223 R | *Il2rb* | CCCATTTCAAAGGGAAGACA |
| 3224 R | *Il2rb* | ACTCTTCCAGGTGAGCTTTG |
| 3225 F | *Il9r* | TTCGAGTCCAGCCTCTTCTA |
| 3227 R | *Il9r* | GACAGTAGCCTCTCTGGTTAATG |
| 3258 F | *Il9r* | GGAAATGGAAGGCAAGATTTAGAG |
| 3259 R | *Il9r* | GCTCCAGGGCAAGATTGATA |
| 3228 F | *Il21r* | GCCCGGATGAAATGTAGTCTTA |
| 3229 R | *Il21r* | GTCGCAAACAGAAAGGGAAATC |
| 3230 R | *Il21r* | ACAACTGAAGTCTCCACAAAGA |
| 3457 F | *Il4* | CTGCCTCCATCATCCTTCTATG |
| 3458 R | *Il4* | GCCAATCAGCACCTCTCTT |
| 3461 R | *Il4* | GAAGAGCAACAGGTACTCCTAAC |
| ***Recombinant DNA*** | N/A |  |
| ***Software and Algorithms*** |  |  |
| Sequencher 4.9 (sequence assembly) | Gene Codes Corporation | https://www.genecodes.com/ |
| GraphPad Prism 10 | GraphPad Software | https://www.graphpad.com; RRID: SCR_002798 |
| FlowJo | BD Biosciences | https://www.flowjo.com; RRID: SCR_008520 |
| R (statistical computing) | R Core Team | https://www.r-project.org; RRID: SCR_001905 |
| DESeq2 (RNA-seq differential expression) | (Love et al., 2014) | https://bioconductor.org/packages/DESeq2; RRID: SCR_015687 |
| GSEA (gene set enrichment analysis) | (Subramanian et al., 2005) | https://www.gsea-msigdb.org; RRID: SCR_003199 |
| GSEApy (Python GSEA library) | (Fang et al., 2023) | https://github.com/zqfang/GSEApy |
| scipy.stats (Python statistical functions) | (Virtanen et al., 2020) | https://scipy.org |
| pandas (Python data analysis) | (McKinney, 2010) | https://pandas.pydata.org |
| seaborn / matplotlib (Python visualization) | (Waskom, 2021)(Hunter, 2007) | https://seaborn.pydata.org; https://matplotlib.org |
| IDT PrimerQuest | IDT | https://www.idtdna.com |
| MfePrimer3 (primer design) | (Wang et al., 2019) | https://mfeprimer3.igenetech.com/spec |
| ***Other*** |  |  |
| NanoDrop ND-1000 UV spectrophotometer | NanoDrop Technologies |  |
| iBLOT Gel Transfer System | Invitrogen/Thermo Fisher | N/A |

**References**

Fang, Z., Liu, X., and Peltz, G. (2023). GSEApy: a comprehensive package for performing gene set enrichment analysis in Python. Bioinformatics *39*.

Grabowski, P., Hesse, S., Hollizeck, S., Rohlfs, M., Behrends, U., Sherkat, R., Tamary, H., Ünal, E., Somech, R., Patıroğlu, T.*, et al.* (2019). Proteome Analysis of Human Neutrophil Granulocytes From Patients With Monogenic Disease Using Data-independent Acquisition. Mol Cell Proteomics *18*, 760-772.

Hunter, J.D. (2007). Matplotlib: A 2D Graphics Environment. Computing in Science & Engineering *9*, 90-95.

Love, M.I., Huber, W., and Anders, S. (2014). Moderated estimation of fold change and dispersion for RNA-seq data with DESeq2. Genome Biol *15*, 550.

Penno, C.A., Jäger, P., Laguerre, C., Hasler, F., Hofmann, A., Gass, S.K., Wettstein-Ling, B., Schaefer, D.J., Avrameas, A., Raulf, F.*, et al.* (2020). Lipidomics Profiling of Hidradenitis Suppurativa Skin Lesions Reveals Lipoxygenase Pathway Dysregulation and Accumulation of Proinflammatory Leukotriene B4. J Invest Dermatol *140*, 2421-2432.e2410.

Rieckmann, J.C., Geiger, R., Hornburg, D., Wolf, T., Kveler, K., Jarrossay, D., Sallusto, F., Shen-Orr, S.S., Lanzavecchia, A., Mann, M.*, et al.* (2017). Social network architecture of human immune cells unveiled by quantitative proteomics. Nat Immunol *18*, 583-593.

Sollberger, G., Brenes, A.J., Warner, J., Arthur, J.S.C., and Howden, A.J.M. (2024). Quantitative proteomics reveals tissue-specific, infection-induced and species-specific neutrophil protein signatures. Sci Rep *14*, 5966.

Subramanian, A., Tamayo, P., Mootha, V.K., Mukherjee, S., Ebert, B.L., Gillette, M.A., Paulovich, A., Pomeroy, S.L., Golub, T.R., Lander, E.S.*, et al.* (2005). Gene set enrichment analysis: a knowledge-based approach for interpreting genome-wide expression profiles. Proc Natl Acad Sci U S A *102*, 15545-15550.

Ter Haar, N.M., Tak, T., Mokry, M., Scholman, R.C., Meerding, J.M., de Jager, W., Verwoerd, A., Foell, D., Vogl, T., Roth, J.*, et al.* (2018). Reversal of Sepsis-Like Features of Neutrophils by Interleukin-1 Blockade in Patients With Systemic-Onset Juvenile Idiopathic Arthritis. Arthritis Rheumatol *70*, 943-956.

Thimmappa, P.Y., Nair, A.S., Najar, M.A., Mohanty, V., Shastry, S., Prasad, T.S.K., and Joshi, M.B. (2022). Quantitative phosphoproteomics reveals diverse stimuli activate distinct signaling pathways during neutrophil activation. Cell Tissue Res *389*, 241-257.

Virtanen, P., Gommers, R., Oliphant, T.E., Haberland, M., Reddy, T., Cournapeau, D., Burovski, E., Peterson, P., Weckesser, W., Bright, J.*, et al.* (2020). SciPy 1.0: fundamental algorithms for scientific computing in Python. Nat Methods *17*, 261-272.

Wang, K., Li, H., Xu, Y., Shao, Q., Yi, J., Wang, R., Cai, W., Hang, X., Zhang, C., Cai, H.*, et al.* (2019). MFEprimer-3.0: quality control for PCR primers. Nucleic Acids Res *47*, W610-w613.
